# Structural and mechanistic basis of reiterative transcription initiation

**DOI:** 10.1101/2021.05.08.443277

**Authors:** Yu Liu, Jared T. Winkelman, Libing Yu, Chirangini Pukhrambam, Emre Firlar, Jason T Kaelber, Yu Zhang, Bryce E. Nickels, Richard H. Ebright

## Abstract

Reiterative transcription initiation, observed at promoters that contain homopolymeric sequences at the transcription start site, generates RNA products having 5’ sequences non-complementary to the DNA template. Here, using crystallography and cryo-EM to define structures, protein-DNA-photocrosslinking to map positions of RNAP leading and trailing edges relative to DNA, and single-molecule DNA nanomanipulation to assess RNAP-dependent DNA unwinding, we show that RNA extension in reiterative transcription initiation: (1) occurs without DNA scrunching, (2) involves a short, 2-3 bp RNA-DNA hybrid, and (3) generates RNA that exits RNAP through the portal by which scrunched nontemplate-strand DNA exits RNAP in standard transcription initiation. The results establish that, whereas RNA extension in standard transcription initiation proceeds through a scrunching mechanism, RNA extension in reiterative transcription initiation proceeds through a slippage mechanism, with slipping of RNA relative to DNA within a short RNA-DNA hybrid, and with extrusion of RNA from RNAP through an alternative RNA exit.

## Introduction

In standard transcription initiation, RNA polymerase (RNAP) holoenzyme binds to promoter DNA, unwinds ∼13 base pairs (bp) of promoter DNA to form an RNAP-promoter open complex (RPo) containing a single-stranded “transcription bubble,” selects a transcription start site, and synthesizes the first 10 nucleotides of the RNA product as an RNAP-promoter initial transcribing complex (RPitc) (Chen et al., 2021; Mazumder and Kapanidis, 2019; Ruff et al., 2015; Winkelman et al., 2021). Standard transcription initiation proceeds through a “DNA scrunching” mechanism, in which RNAP unwinds additional DNA, pulls the additional unwound DNA past its active center, and accommodates the additional unwound DNA as single-stranded bulges within the transcription bubble (Kapanidis et al., 2006; Margeat et al., 2006; Revyakin et al., 2006; Winkelman et al., 2015). During standard transcription initiation, each step of RNA extension involves: (i) unwinding of 1 bp of DNA downstream of the RNAP active center, expanding the transcription bubble by 1 bp; (ii) translocation of DNA and RNA--together--by 1 bp relative to the RNAP active center by 1 bp; (iii) binding, through base pairing, of a complementary nucleoside triphosphate (NTP) to the DNA template strand in the RNAP active center, and (iv) phosphodiester bond formation, resulting in addition of a nucleotide to the RNA 3’ end (Winkelman et al., 2021). Standard transcription initiation yields an RNA product having a sequence fully complementary to the DNA template strand. Furthermore, during standard transcription initiation, the RNA product remains fully base paired to the DNA template strand, as an RNA-DNA “hybrid.”

In an alternative pathway of transcription initiation, termed “reiterative transcription initiation,” “transcriptional stuttering,” or “pseudo-templated transcription,” an RNAP-promoter reiteratively transcribing complex (RPrtc) synthesizes an RNA product having a 5’-end sequence that contains a variable number, up to tens to hundreds, of nucleotides not complementary to the DNA template (Jacques and Kolakofsky, 1991; Turnbough and Switzer, 2008; Turnbough Jr, 2011). Reiterative transcription initiation, which was first observed six decades ago (Chamberlin and Berg, 1962, 1964), competes with standard transcription initiation, both *in vitro* and *in vivo* (Cheng et al., 2001; Guo and Roberts, 1990; Han and Turnbough, 1998; Jensen-MacAllister et al., 2007; Liu et al., 1994; Meng et al., 2004; Qi and Turnbough, 1995; Severinov and Goldfarb, 1994; Tu and Turnbough, 1997; Vvedenskaya et al., 2015; Wagner et al., 1990). Reiterative transcription initiation is observed at promoters that contain homopolymeric sequences at, or immediately downstream of, the transcription start site, resulting in low yields of standard, full-length RNA products at such promoters (Jacques and Kolakofsky, 1991; Turnbough and Switzer, 2008; Turnbough Jr, 2011). The extent of reiterative transcription initiation relative to standard initiation can change in response to changes in nucleoside triphosphate (NTP) concentrations (Turnbough, 2019; Turnbough and Switzer, 2008; Turnbough Jr, 2011). Classic examples of genes regulated through changes in the extent of reiterative transcription initiation relative to standard transcription initiation in response to changes in NTP concentrations are *Escherichia coli pyrBI* and *Bacillus subtilis pyrG* (Liu et al., 1994; Meng et al., 2004).

The mechanism of reiterative transcription initiation has not been firmly established. It has been hypothesized that reiterative transcription initiation involves an “RNA slipping” mechanism, in which RNA extension does not involve translocation of DNA relative to the RNAP active center, but, instead, involves translocation of RNA--“slippage”--relative to both DNA and the RNAP active center (Guo and Roberts, 1990; Jacques and Kolakofsky, 1991; Macdonald et al., 1993; Murakami et al., 2017; Severinov and Goldfarb, 1994; Shin et al., 2020; Turnbough and Switzer, 2008; Turnbough Jr, 2011; Vvedenskaya et al., 2015; Wagner et al., 1990). However, direct evidence for RNA slipping has not been presented, and the mechanism by which long reiteratively transcribed RNA products--RNA products tens to hundreds of nucleotides in length--leave the RNAP active center has not been defined.

Crystal structures have been reported of RPrtc (Murakami et al., 2017; Shin et al., 2020). However, in those structures, (i) transcription-bubble nontemplate-strand DNA was extensively disordered (3-8 nt disordered), complicating assessment of nontemplate-strand DNA scrunching; (ii) transcription-bubble template-strand DNA was partly missing and partly disordered (5 nt missing; 3 nt disordered), precluding assessment of template-strand DNA scrunching; (iii) the position of the RNA 3’-end relative to the DNA template strand did not permit further reiterative transcription (template-strand homopolymeric sequence not aligned with RNAP active-center addition site), precluding assessment of the mechanism of RNA extension; and (iv) only complexes with short RNA products (6-8 nt) were analyzed, precluding assessment of how long reiterative-transcription-generated RNA products exit the RNAP active-center cleft (Murakami et al., 2017; Shin et al., 2020). As a result of these limitations, the previous crystal structures did not enable determination of the roles of DNA scrunching and RNA slipping, the length of the RNA-DNA hybrid, and the RNA exit path in reiterative transcription initiation

Here, we report crystal structures of RNAP engaged in standard transcription initiation of short RNA products on a template containing a non-homopolymeric sequence and RNAP engaged in reiterative transcription initiation of short RNA products on templates containing template-strand GGG and CCC homopolymeric sequences. In addition, we report a cryo-EM structure of RNAP engaged in reiterative transcription initiation of long--up to at least 50 nt--RNA products on a template containing a template-strand CCC homopolymeric sequence. The structures reveal that, whereas RNA extension in standard transcription initiation involves DNA scrunching, RNA extension in reiterative transcription initiation does not. The structures further reveal that only two template-strand nucleotides (in a post-translocated state) or three template-strand nucleotides (in a pre-translocated state) are positioned to be able to base pair to the RNA product, resulting in a short RNA-DNA hybrid. The cryo-EM structure of RNAP engaged in reiterative transcription initiation of long RNA products further reveals that reiterative-transcription-generated RNA exits RNAP using the path by which scrunched nontemplate-strand DNA exits RNAP in standard transcription initiation, instead of the path by which RNA exits RNAP in standard transcription initiation. Results of two independent orthogonal approaches, site-specific protein-DNA photocrosslinking and single-molecule DNA nanomanipulation, confirm the observed scrunching patterns. Taken together, our results establish that, whereas RNA extension in standard initiation involves DNA scrunching, RNA extension in reiterative initiation involves RNA slipping, with sliding of the RNA product relative to the DNA template strand within a short RNA-DNA hybrid, and with extrusion of RNA from the RNAP active-center cleft through an alternative RNA exit.

## Results

### Crystal structures of RPrtc,4 and RPrtc,5: RNA extension through RNA slipping without DNA scrunching

We determined crystal structures of *Thermus thermophilus* RPrtc in which nontemplate-strand DNA is fully ordered, enabling assessment of DNA scrunching, and in which the template-strand homopolymeric sequence is aligned with the RNAP active-center addition site, enabling assessment of RNA slipping (Figures 1-2). For both the crystal structures of this section and the cryo-EM structure below, we analyzed *T. thermophilus* RNAP because atoms of this hyperthermophilic bacterial RNAP show lower thermal motions at structure-determination temperatures than atoms of mesophilic bacterial RNAP, enabling determination of structures having high order, higher resolution, and superior map quality (Basu et al., 2014; Feng et al., 2016; Murakami et al., 2002a; Murakami et al., 2002b; Murakami et al., 2017; Shin et al., 2020; Vassylyev et al., 2002; Zhang et al., 1999; Zhang et al., 2012). We obtained a crystal structure of RPrtc containing a 5 nt RNA product by incubation of a nucleic-acid scaffold containing a G_+1_G_+2_G_+3_ template-strand homopolymeric sequence, where +1 is the transcription start site, with RNAP and CTP (RPrtc,5 [G_+1_G_+2_G_+3_]; Figures 1C and S1A; Table S1), and we obtained a crystal structure of RPrtc containing a 4 nt RNA product by incubation of a nucleic-acid scaffold containing a C_+1_C_+2_C_+3_ template-strand homopolymeric sequence with RNAP and GTP (RPrtc,4 [C_+1_C_+2_C_+3_]; Figures 1D and S1A; Table S1). For reference, we compared these structures to a previously reported crystal structure of *T. thermophilus* RPo (RPo; Figure 1A, Zhang et al., 2012) and a crystal structure of *T. thermophilus* RPitc containing a 5 nt RNA product by incubation of a nucleic-acid scaffold lacking a template-strand homopolymeric sequence with RNAP, ATP, UTP, and CTP (RPitc,5; Figures 1B and S1A; Table S1).

**Figure 1.**
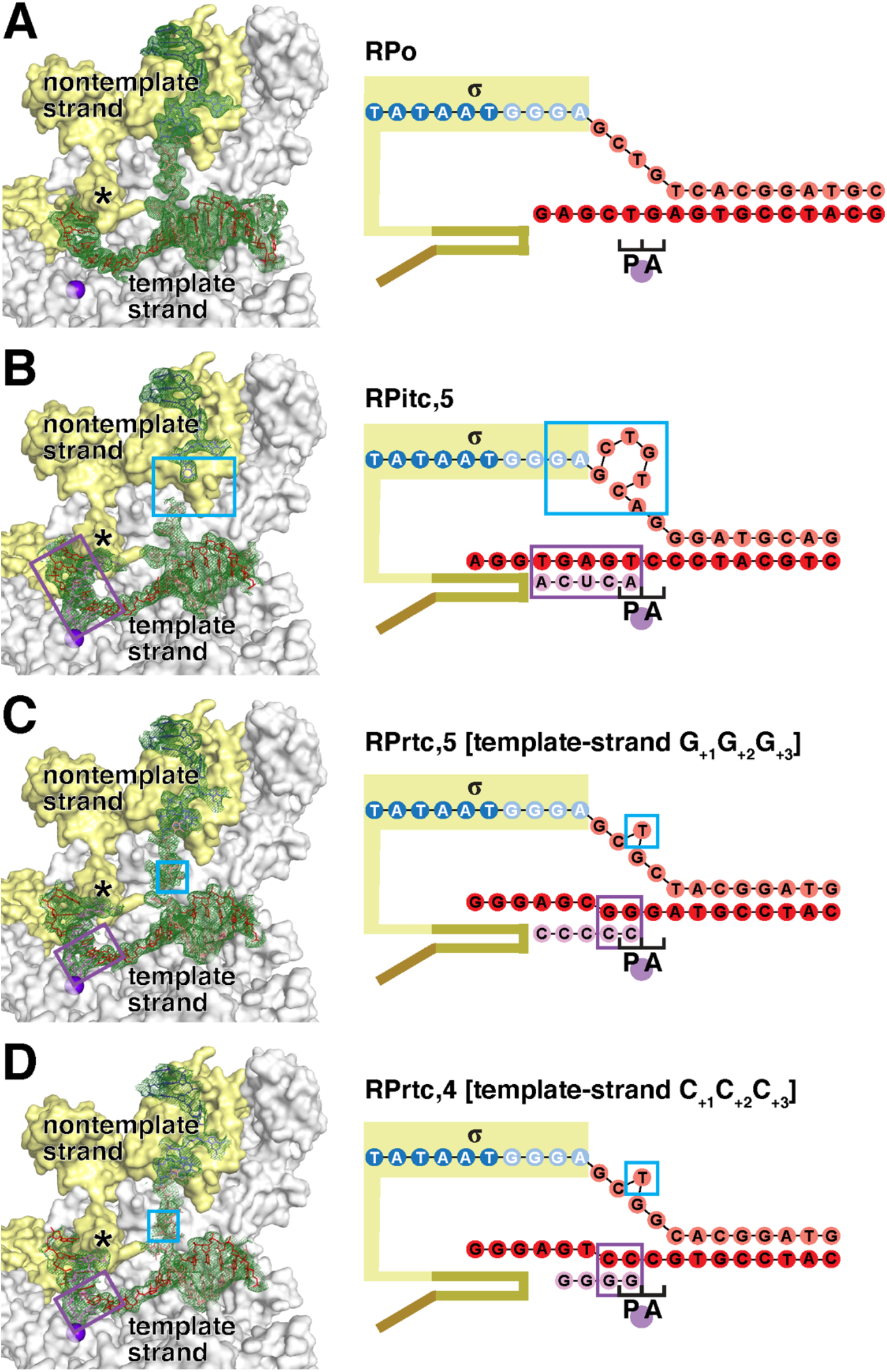
Crystal structures of RPrtc,4 and RPrtc,5: RNA extension through RNA slipping without DNA scrunching. Crystal structures of transcription initiation complexes engaged in standard transcription initiation and reiterative transcription initiation. Left, experimental electron density (mFo-DFc; contoured at 2.0σ in A and 1.5σ in B-D) and atomic model, showing interactions of RNAP and σ with transcription-bubble nontemplate strand, transcription-bubble template strand, and downstream dsDNA (RNAP β subunit and β’ non-conserved domain omitted for clarity). Right, nucleic-acid scaffold. RNAP, gray; RNAP active-center catalytic Mg^2+^(I) ion, violet filled circle; σ, yellow; σ finger, asterisk (left subpanel) and yellow-brown (right subpanel); σR3-σR4 linker in RNA exit channel, brown; −10 element of DNA nontemplate strand, dark blue; discriminator element of DNA nontemplate strand, light blue; rest of DNA nontemplate strand, pink; DNA template strand, red; RNA product, magenta. Cyan rectangles on panel B indicate disordered regions containing scrunched nucleotides. Cyan rectangles in panels C and D indicate ordered scrunched nucleotides. Bulged-out nucleotides in panels B-D, right, indicate bulged-out scrunched nucleotides. Violet rectangles indicate RNA-DNA hybrids. Raised template-strand nucleotides in panels C-D indicate non-base-paired nucleotides. **(A)** RNAP-promoter open complex (RPo; PDB 4G7H; Zhang et al., 2012). **(B)** RNAP-promoter initial transcribing complex containing 5 nt RNA product generated by *in crystallo* standard transcription initiation (RPitc,5). **(C)** RNAP-promoter reiteratively transcribing complex containing 5 nt RNA product generated by *in crystallo* reiterative transcription initiation on nucleic-acid scaffold having a template-strand G_+1_G_+2_G_+3_ homopolymeric sequence (RPrtc,5 [G_+1_G_+2_G_+3_]). **(D)** RNAP-promoter reiteratively transcribing complex containing 4 nt RNA product generated by *in crystallo* reiterative transcription initiation on nucleic-acid scaffold having a template-strand C_+1_C_+2_C_+3_ homopolymeric sequence (RPrtc,4 [C_+1_C_+2_C_+3_]).

**Figure 2.**
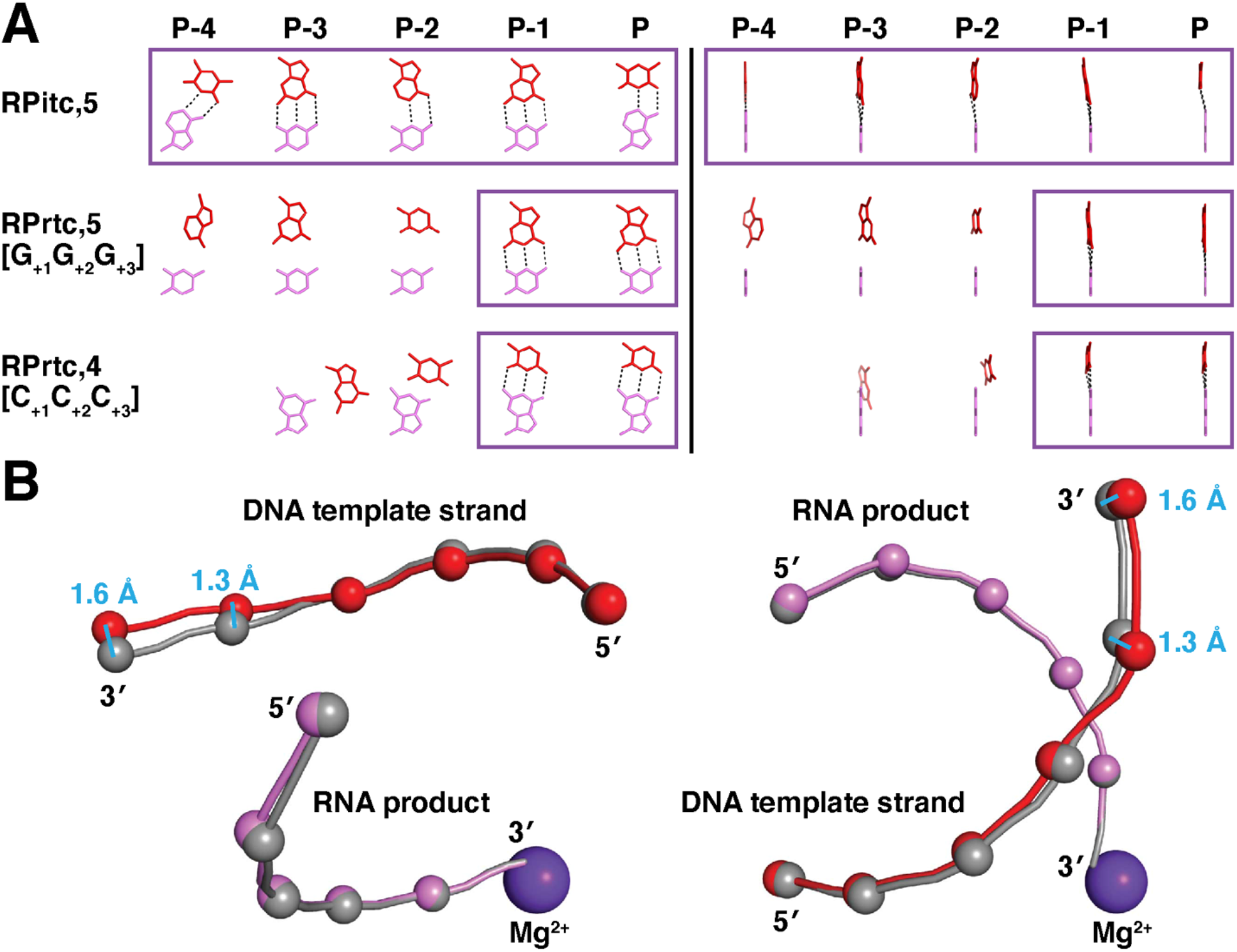
Crystal structures of RPrtc,4 and RPrtc,5: short RNA-DNA hybrid. **(A)** RNA-DNA base pairing in crystal structures of transcription initiation complexes engaged in standard transcription initiation (RPitc,5) and reiterative transcription initiation (RPrtc,5 [G_+1_G_+2_G_+3_] and RPrtc,4 [C_+1_C_+2_C_+3_]). Left, template-strand DNA bases (red) and corresponding RNA bases (magenta) in view orientation parallel to RNA-DNA-hybrid helix axis. Right, template-strand DNA bases (red) and corresponding RNA bases (magenta) in view orientation perpendicular to RNA-DNA-hybrid helix axis. Positions are numbered relative to the RNAP active-center P site. Dashed lines indicate Watson-Crick H-bonds. Violet rectangles indicate RNA-DNA hybrids. At positions P-4, P-3, and P-2 of RPrtc,5 [G_+1_G_+2_G_+3_], and at positions P-3 and P-2 of RPrtc,4 [C_+1_C_+2_C_+3_], template-strand DNA bases are displaced relative to their locations in RPitc,5, and no base pairing occurs. **(B)** Superimposition of DNA template strand and RNA of RPrtc,5 [G_+1_G_+2_G_+3_] (red spheres, DNA phosphates; magenta spheres, RNA phosphates; violet sphere, RNAP active-center catalytic Mg^2+^ ion) on DNA template strand and RNA of RPitc,5 (gray spheres, DNA and RNA phosphates). Left, view orientation parallel to RNA-DNA-hybrid helix axis; right, view orientation perpendicular to RNA-DNA-hybrid helix axis. Distances in cyan, displacement of template-strand DNA nucleotides at positions P-4 and P-3 of RPrtc,5 [G_+1_G_+2_G_+3_] relative to their locations in RPitc,5.

The crystal structure of RPo shows ordered density for all nucleotides of the transcription-bubble nontemplate strand: 5 nt in the −10 element (promoter element recognized by conserved region 2 of transcription initiation factor σ; Feklistov et al., 2014), 4 nt in the discriminator element (another promoter element recognized by conserved region 2 of transcription initiation factor σ; Feklistov et al., 2014), and 4 nt between the discriminator element and downstream dsDNA; Figure 1A, blue, light blue, and pink; (Zhang et al., 2012).

The crystal structure of RPitc,5 shows an initially transcribing complex with a 5 nt RNA product and an RNAP active-center post-translocated state (Figure 1B). The RNA product is fully base paired to the DNA template strand as an RNA-DNA hybrid, with the RNA 3’ nucleotide and the corresponding DNA template-strand nucleotide located in the RNAP active-center product site (“P site”), and the next DNA template-strand nucleotide in the RNAP active-center addition site (“A site”) available for base pairing with an incoming NTP. The positions of the RNA and DNA relative to the RNAP active center indicate that, as compared to in RPo, 4 bp of downstream dsDNA have been unwound, 4 nt of each strand has been translocated relative to the RNAP active center, and the RNA product has translocated 4 nt in lock-step register with template-strand DNA. The crystal structure of RPitc,5 indicates that the 4 nt of nontemplate-strand DNA translocated relative to the RNAP active center are accommodated through DNA scrunching, with bulging of the nontemplate-strand DNA segment between the discriminator element and downstream dsDNA. Thus, the crystal structure of RPitc,5 shows ordered density for all transcription-bubble nontemplate-strand nucleotides of the −10 element and the upstream half of the discriminator element, with the same positions and the same σ-DNA interactions as in RPo, and shows disorder for 8 nt of nontemplate-strand DNA, corresponding to the downstream half of the discriminator element and DNA immediately downstream of the discriminator element (Figure 1B, cyan boxes). The 8 nt segment of disordered nontemplate-strand DNA in RPitc,5 has exactly the same endpoints, and spans exactly the same distance, as a 4 nt segment of ordered nontemplate-strand DNA in RPo (Figures 1A-B), indicating that ∼4 nt of the 8 nt segment of disordered nontemplate-strand DNA are flipped out and/or bulged out relative to the path of the nontemplate strand in RPo (Figure 1B, right, flipped and/or bulged nucleotides in cyan box). The disorder of the 8 nt segment of disordered nontemplate-strand DNA indicates that the segment adopts an ensemble of distinct conformations (Figure 1B, cyan boxes). We conclude that formation of RPitc,5 involves ∼4 bp of DNA scrunching.

The crystal structure of RPrtc,5 [G_+1_G_+2_G_+3_] shows a reiteratively transcribing complex with a 5 nt RNA product and an RNAP active-center post-translocated state (Figure 1C). Only part of the RNA product--the part comprising the 3’ nucleotide and the adjacent nucleotide--is base paired to the DNA template strand as a 2 bp RNA-DNA hybrid, with the RNA 3’ nucleotide and the corresponding DNA template-strand nucleotide located in the RNAP active-center P site, and the next DNA template-strand nucleotide in the RNAP active-center A site available for base pairing with an incoming NTP. The positions of the RNA and DNA relative to the RNAP active center indicate that, as compared to in RPo, 1 bp of downstream dsDNA has been unwound, 1 nt of each strand of DNA has been translocated relative to the RNAP active center, and the 5’ end of the RNA product has been translocated by 4 nt relative to the RNAP active center, translocating 1 nt in register with template-strand DNA and 3 nt out of register with template-strand DNA. The crystal structure of RPrtc,5 [G_+1_G_+2_G_+3_] shows that the 1 nt of nontemplate-strand DNA translocated relative to the RNAP active center is accommodated through DNA scrunching, with unstacking and flipping of 1 nt of nontemplate-strand DNA between the discriminator element and downstream dsDNA (Figure 1C, cyan boxes). The crystal structure of RPrtc,5 [G_+1_G_+2_G_+3_] shows ordered density for all transcription-bubble nontemplate-strand nucleotides, including the scrunched, unstacked, flipped nucleotide: 5 nt in the −10 element, 4 nt in the discriminator element, and 5 nt between the discriminator element and downstream dsDNA (Figure 1C). The crystal structure shows graphically that formation of RPrtc,5 [G_+1_G_+2_G_+3_] involves 1 bp of DNA scrunching and 3 nt of RNA slipping.

The crystal structure of RPrtc,4 [C_+1_C_+2_C_+3_] shows a reiteratively transcribing complex with a 4 nt RNA product and an RNAP active-center post-translocated state, a 2 bp RNA-DNA hybrid, 1 bp of unwinding of downstream dsDNA, 1 nt of translocation of each DNA strand, 1 nt of translocation of the 5’ end of the RNA product in register with template-strand DNA, and 2 nt of translocation of the 5’ end of the RNA product out of register with template-strand DNA (Figure 1D). The crystal structure shows graphically that formation of RPrtc,4 [C_+1_C_+2_C_+3_] involves 1 bp of DNA scrunching and 2 nt of RNA slipping.

We infer, based on the structures in Figure 1, that standard transcription initiation to generate a post-translocated state of RPitc,x involves x-1 bp of DNA scrunching, whereas reiterative transcription initiation to form a post-translocated state of RPrtc,x involves 1 bp of DNA scrunching and x-2 nt of RNA slipping. Expressing these inferences in terms of mechanism, we infer that, in standard transcription initiation, following synthesis of a 2 nt initial RNA product, 1 bp of DNA scrunching occurs for each 1 nt of RNA extension (Revyakin et al., 2006), and we infer that, in contrast, in reiterative transcription initiation, following synthesis of a 2 nt initial RNA product, 1 bp of DNA scrunching occurs to position the 3’ end of the initial RNA product in the RNAP active-center P site, and no further DNA scrunching--just RNA slipping--occurs in RNA extension (see Discussion).

### Crystal structures of RPrtc,4 and RPrtc,5: short RNA-DNA hybrid

In the crystal structure of RPitc,5, all nucleotides of the RNA product are complementary to, and base-paired to, the DNA template strand, yielding a 5 bp RNA-DNA hybrid (Figures 1B and 2A). In contrast, in the crystal structures of RPrtc,5 [G_+1_G_+2_G_+3_] and RPrtc,4 [C_+1_C_+2_C_+3_], only the 2 nt at the 3’ end of the RNA product are complementary to, and base-paired to, the DNA template strand, yielding a 2 bp RNA-DNA hybrid (Figures 1C-D and 2A). We conclude that, unlike standard transcription initiation, reiterative transcription initiation involves a short RNA-DNA hybrid: a hybrid that is only 2 bp in length in the RNAP active-center post-translocated state of the crystal structures (“post-slipped state”), and that would be only 3 bp in length upon NTP binding and phosphodiester bond formation to yield the RNAP active-center pre-translocated state (“pre-slipped state”).

In RPrtc, the conformation and interactions with RNAP of the RNA product--even the part of the RNA product that is not complementary to, and not base paired to, template-strand DNA--are the same as in RPitc (Figure 2B). In contrast, in RPrtc, the conformation and interactions of the part of the DNA template strand that is not complementary to, and not base paired to, RNA differ from those in RPitc (Figure 2B). Inspection of structures of RPrtc and RPitc indicates that RNAP makes numerous interactions with RNA, and few interactions with DNA, in the RNAP hybrid-binding region, accounting for our observation that RNA conformation, rather than DNA conformation, is maintained upon loss of RNA-DNA complementarity and base pairing.

In our crystal structures of RPitc and RPrtc, the 5’ end of the RNA product is in contact with the “σ finger” (also referred to as σ region 3.2), which enters the RNAP active-center cleft, and obstructs the path of the RNA product (Figure 1; see, Basu et al., 2014; Li et al., 2020; Mazumder and Kapanidis, 2019; Mekler et al., 2002; Murakami et al., 2002b; Vassylyev et al., 2002; Winkelman et al., 2021; Zhang et al., 2012). In solution, and in some crystal forms under some conditions, extension of the RNA product beyond a length of 5 nt can drive stepwise displacement of the σ finger (Li et al., 2020). In contrast, with the crystal forms and conditions of this work, RNA extension does not drive stepwise displacement of the σ finger--presumably due to crystal-lattice constraints on conformational change in σ--and thus RNA products are limited to a length of 5 nt.

### Cryo-EM structure of RPrtc,≥11: RNA extension through RNA slipping without DNA scrunching

Reiterative transcription initiation can generate RNA products up to at least 50 nt in length (Figure S1; Turnbough and Switzer, 2008; Turnbough Jr, 2011). Because the volume of the RNAP active center cleft in RPrtc cannot accommodate more than ∼10 nt of RNA, long RNA products generated by reiterative transcription initiation must exit from, and extend outside, the RNAP active-center cleft (Murakami et al., 2017). A key unresolved question is where long RNA products generated in reiterative transcription initiation exit the RNAP active-center cleft. In the crystal structures of this work, as in the crystal structures of Murakami et al., 2017 and Shin et al., 2020, RNA products generated by reiterative transcription initiation were limited in length, because further RNA extension was blocked by the presence of the σ finger in the RNAP active-center cleft and by crystal-lattice constraints that prevented displacement of the σ finger from the RNAP active-center cleft (Basu et al., 2014; Li et al., 2020), opening of the RNAP clamp (Chakraborty et al., 2012; Duchi et al., 2018; Mazumder et al., 2020), or any other conformational change that could open a path for further extension of RNA and for extrusion of RNA from the RNAP active-center cleft. One hypothesis is that, in solution, complete displacement of the σ finger from the RNAP active-center cleft channel could allow long RNA products generated in reiterative transcription initiation to exit the RNAP active-center cleft through the RNAP RNA-exit channel, the same exit route used by RNA in standard transcription (Murakami et al., 2017). Another hypothesis is that, in solution, a smaller conformational change in σ and/or RNAP could allow long RNA products generated in reiterative transcription initiation to exit the RNAP active-center cleft through a different route (Murakami et al., 2017). Consistent with the first hypothesis, our crystal structures show that the 3’ region of RNA products generated in reiterative transcription can follow the same path relative to RNAP as in standard transcription initiation (Figure 2). However, arguing against the first hypothesis, it is unclear how, with only 1 bp of DNA scrunching (Figure 1), the system could acquire the energy needed to drive complete displacement of the σ finger from the RNAP hybrid binding site and displacement of the σ region-3/region-4 linker from the RNAP RNA-exit channel (free energy that, in standard transcription initiation, is thought to be provided by ∼8-10 bp of DNA scrunching; Kapanidis et al., 2006; Revyakin et al., 2006), and it is unclear how the σ region-3/region-4 linker could be completely displaced from the RNAP RNA-exit channel without triggering promoter escape (which, in standard transcription initiation, is thought to be triggered by displacement of the region-3/region-4 linker from the RNAP RNA-exit channel; Li et al., 2020; Mekler et al., 2002; Murakami et al., 2002b; Vassylyev et al., 2002).

To resolve these questions, we performed cryo-EM structure determination, analyzing reiteratively transcribing complexes prepared in solution. We incubated a nucleic-acid scaffold containing a full transcription bubble and a C_+1_C_+2_C_+3_ template-strand homopolymeric sequence with RNAP and GTP, we applied samples to glow-discharged graphene-oxide-coated grids (Pantelic et al., 2010; Thomas et al., 2016), flash-froze samples, and performed single particle-reconstruction cryo-EM (RPrtc,≥11 [C_+1_C_+2_C_+3_]; Figures 3-4, S1A, and S2). This approach avoided the limitation imposed by crystal-lattice constraints (Figures 3-4 vs. Figures 1-2 and Murakami et al., 2017; Shin et al., 2020). In addition, by analyzing a nucleic-acid scaffold that contained a full transcription bubble, this approach avoided possible limitations imposed by use of a nucleic-acid scaffold that lacked an upstream duplex (Figures 3-4 vs. Figures 1-2 and Murakami et al., 2017; Shin et al., 2020). Use of glow-discharged graphene-oxide-coated grids was essential in order to obtain a satisfactory distribution of particle orientations on grids (Figure S2D).

**Figure 3.**
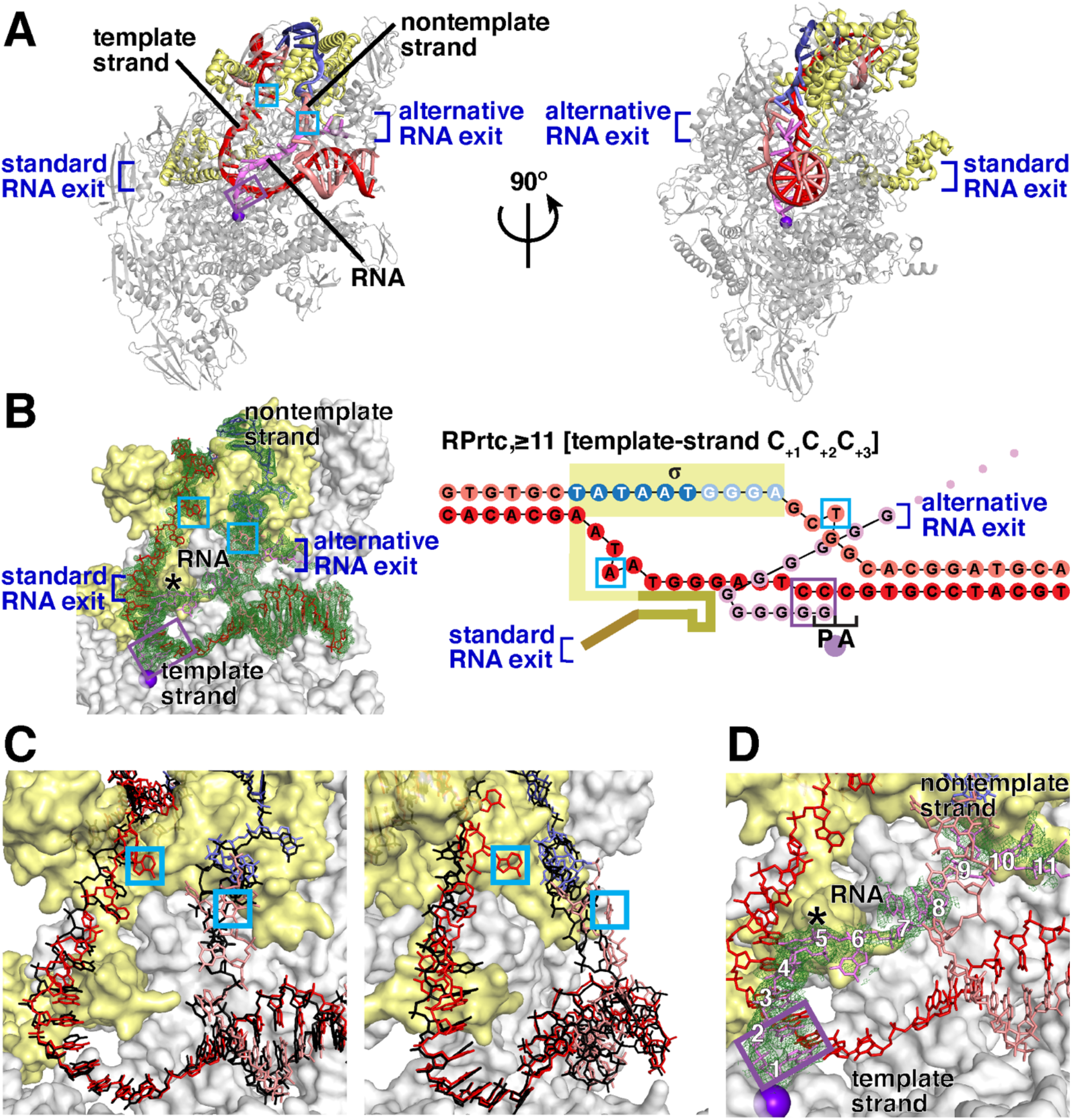
Cryo-EM structure of RPrtc,≥11: RNA extension through RNA slipping without DNA scrunching. **(A)** Overall structure (β’ nonconserved region omitted for clarity; two orthogonal view orientations). Dark blue brackets indicate the standard RNA exit and alternative RNA exit. Cyan rectangles indicate scrunched nucleotides. Violet rectangles indicate RNA-DNA hybrids. Other symbols and colors in panels A-D are as in Figure 1. **(B)** Left, cryo-EM density and atomic model, showing interactions of RNAP and σ with transcription-bubble nontemplate strand, transcription-bubble template strand, and downstream dsDNA. Right, nucleic-acid scaffold. Yellow-brown, σ finger (note displacement of σ-finger tip); magenta dots, RNA outside RNAP active-center cleft (nucleotides rN≥11). **(C)** Superimposition of DNA in RPrtc,≥11 [C_+1_C_+2_C_+3_] (pink and red) on DNA in RPo (black; PDB 512D; Feng et al., 2016). **(D)** Close-up of cryo-EM density and atomic model for RNA (nucleotides rN1-rN11 numbered in white).

**Figure 4.**
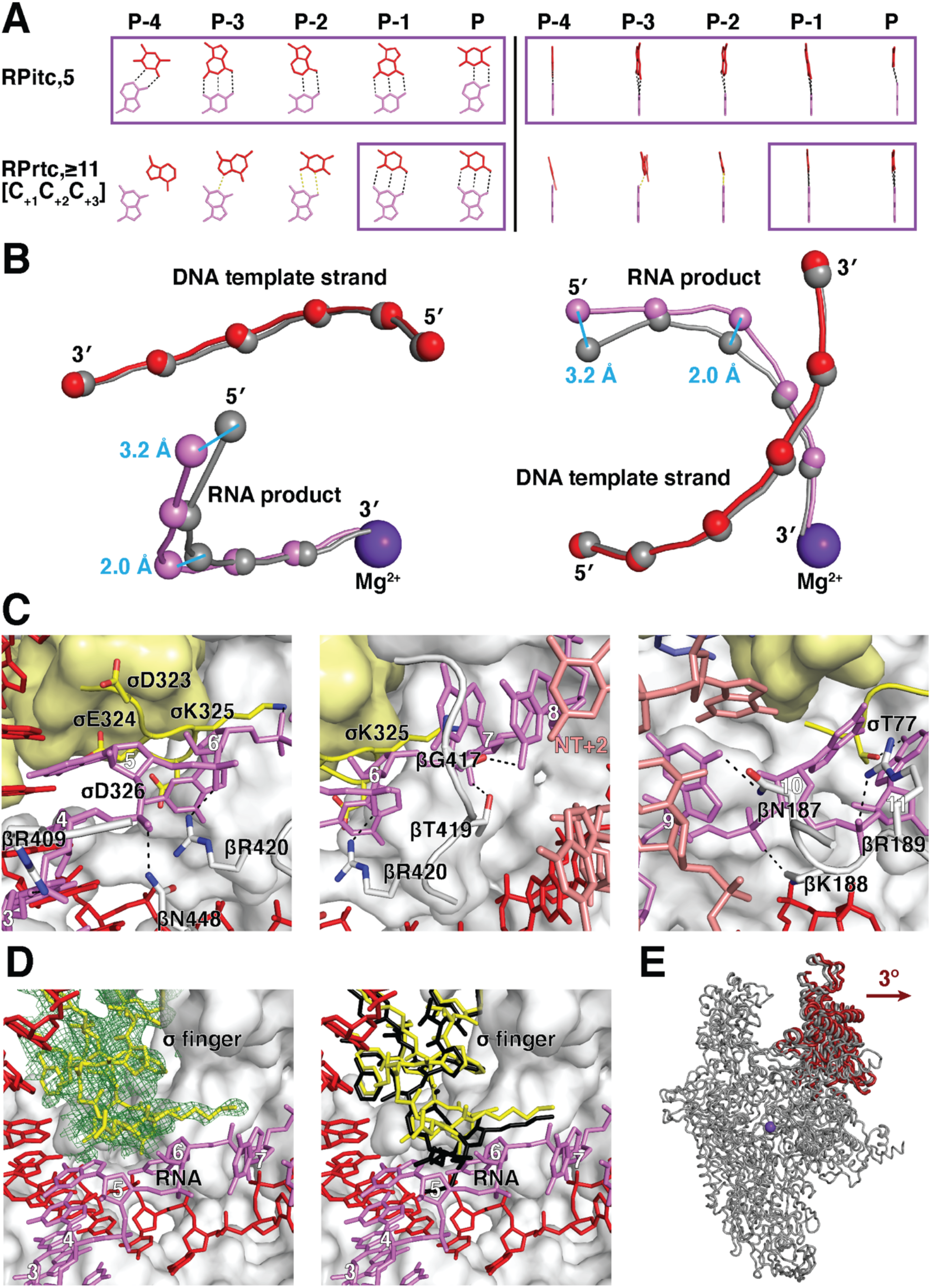
Cryo-EM structure of RPrtc,≥11: short RNA-DNA hybrid and RNA exit through nontemplate-strand scrunching portal. **(A)** RNA-DNA base pairing. Yellow dashed lines, non-Watson-Crick H-bonds. Other symbols and colors as in Figure 2A. **(B)** Superimposition of DNA template strand and RNA of RPrtc,≥11 [C_+1_C_+2_C_+3_] (red spheres, DNA phosphates; magenta spheres, RNA phosphates; violet sphere, RNAP active-center catalytic Mg^2+^ ion) on DNA template strand and RNA of RPitc,5 (gray spheres, DNA and RNA phosphates). **(C)** Protein-RNA and DNA-RNA interactions in alternative RNA path. Left, center, and right subpanels show interactions with RNA nucleotides with rN3-rN6, rN6-rN8, and rN9-rN11 (atoms in magenta; nucleotide numbers in white). Ribbons and sticks, backbone segments and sidechain atoms of residues of RNAP β (backbone segments in white; carbon oxygen and nitrogen atoms in white, red, and blue) and σ (backbone segments in yellow; carbon oxygen and nitrogen atoms in yellow, red, and blue) that make protein-RNA interactions with rN3-rN11. “NT+2,” nucleotide at nontemplate-strand position +2 that makes base-stacking interaction with RNA nucleotide rN8. Other colors as in Figure 1. **(D)** σ-finger conformation. Left, cryo-EM density and atomic model for σ finger. Right, superimposition of σ finger of RPrtc,≥11 [C_+1_C_+2_C_+3_] (yellow) on σ finger of RPo (black; PDB 4G7H; Zhang et al., 2012). Other colors as in Figure 1. **(E)** RNAP clamp conformation. Superimposition of RNAP of RPrtc,≥11 [C_+1_C_+2_C_+3_] (clamp in red; rest of RNAP in gray) on RNAP of RPo (black; PDB 4G7H; Zhang et al., 2012).

The cryo-EM structure of RPrtc,≥11 [C_+1_C_+2_C_+3_] has an overall resolution of 3.0 Å, with higher local resolution for regions of interest, including the transcription-bubble nontemplate and template DNA strands and the RNA product (Figure S2D-E). Map quality is high, with ordered, traceable density for 7 bp of upstream dsDNA, all nucleotides of the nontemplate and template strands of the transcription bubble, 10 bp of downstream dsDNA, and 11 nt of the RNA product corresponding to the 11 nt segment containing the RNA 3’ end (Figures 3-4 and S2E).

The cryo-EM structure of RPrtc,≥11 [C_+1_C_+2_C_+3_] shows a reiteratively transcribing complex with a long, ≥11 nt, RNA product and an RNAP active-center post-translocated state (Figure 3A-B). As observed in our crystal structures of RPrtc with short RNA products (Figure 1C-D), the 2 nt at the 3’ end of the RNA product are base paired to the DNA template strand and the next 3 nt of the RNA product are close to, but not Watson-Crick base-paired with, the DNA template strand (Figure 3A-B). The next 6 nt of the RNA product follow a previously unobserved path that diverges from the DNA template strand due to a collision with the σ finger, crosses the transcription bubble, crosses the DNA nontemplate strand, and exits the RNAP active-center cleft at a position on the face of RNAP opposite the standard RNA exit (“alternative RNA exit”; Figure 3A-B). Additional nucleotides of the long, tens to hundreds of nucleotides, RNA products generated by reiterative transcription under these conditions (Figure S1A) would be located outside the alternative RNA exit and would extend into bulk solvent; these nucleotides are not observed in the structure, presumably due to segmental disorder (Figure 3A-B), analogous to the segmental disorder previously observed for nucleotides of long RNA products located outside the RNAP standard RNA exit (Yin et al., 2019). The structure contains σ and exhibits the same σ-DNA and σ-RNAP interactions, except for those made by the σ-finger tip (see below), as in RPo (Figure 3A-B), indicating that production of long RNAs by reiterative transcription initiation does not involve substantial disruption of σ-DNA and σ-RNAP interactions and does not involve promoter escape.

As observed in our crystal structures of RPrtc with short RNA products (Figure 1C-D), the positions of the RNA and DNA relative to the RNAP active center indicate that, as compared to in RPo, 1 bp of downstream dsDNA has been unwound, 1 nt of each strand of DNA has been translocated relative to the RNAP active center, and the 5’ end of the RNA product has been translocated by ≥11 nt relative to the RNAP active center, translocating 1 nt in register with template-strand DNA and ≥10 nt out of register with template-strand DNA. The cryo-EM structure of RPrtc,≥11 [C_+1_C_+2_C_+3_] shows that the 1 nt of nontemplate-strand DNA translocated relative to the RNAP active center is accommodated through 1 bp of DNA scrunching, with unstacking and flipping of nontemplate-strand position +1 (i.e., the position 7 bp downstream of the −10 element) and with changes in conformation of the nucleotides flanking nontemplate-strand position +1 (Figure 3B-C, cyan boxes). The cryo-EM structure of RPrtc,≥11 [C_+1_C_+2_C_+3_] further shows that the 1 nt of template-strand DNA translocated relative to the RNAP active center likewise is accommodated through 1 bp of DNA scrunching, with unstacking and flipping of template-strand position −9 (i.e., the third position within the −10 element) and with small changes in conformation of template-strand nucleotides downstream of position −9 (Figure 3B-C, cyan boxes). The cryo-EM structure shows graphically that formation of RPrtc,≥11 [C_+1_C_+2_C_+3_] involves 1 bp of DNA scrunching and ≥10 nt of RNA slipping.

### Cryo-EM structure of RPrtc,≥11: short RNA-DNA hybrid

As observed in our crystal structures of RPrtc with short RNA products (Figure 2A), in the cryo-EM structure of RPrtc,≥11 [C_+1_C_+2_C_+3_], only the 2 nt at the 3’ end of the RNA product are complementary to, and base paired to, the DNA template strand, yielding a 2 bp RNA-DNA hybrid (Figures 3A-B and 4A). The structure thus supports our conclusion that, unlike standard transcription initiation, reiterative transcription initiation involves a short RNA-DNA hybrid: a 2 bp hybrid in the post-slipped state, and a 3 bp hybrid in the pre-slipped state.

In our crystal structures of RPrtc with short RNA products, the conformation of the RNA nucleotides at positions P-4, P-3, and P-2 relative to the active-center P site--the RNA nucleotides, not complementary to, and not Watson-Crick base-paired to, template-strand DNA--is the same as in a standard transcription initiation complex, and the conformation of the corresponding part of template-strand DNA is different (Figure 2B). In contrast, in the cryo-EM structure of RPrtc,≥11 [C_+1_C_+2_C_+3_], the opposite is true: the conformation of the RNA nucleotides at positions P-4, P-3, and P-2 relative to the active-center P site differs from the conformation in standard transcription initiation and elongation complexes, and the conformation of the corresponding part of template-strand DNA is the same as in standard transcription initiation and elongation complexes (Figure 4B). The conformations of the RNA product and the DNA template strand at positions P-4, P-3, and P-2 in RPrtc,≥11 [C_+1_C_+2_C_+3_] allow formation of non-Watson-Crick, wobble or wobble-like, H-bonds at positions P-3 and P-2 (Figure 4A). The difference in conformation of RNA and DNA at positions P-4, P-3. and P-2 in our crystal structures of RPrtc with short RNA products and our cryo-EM structure of RPrtc,≥11 [C_+1_C_+2_C_+3_] likely is attributable to the absence in the former, and presence in the latter, of additional RNA nucleotides 5’ to this RNA segment and additional template-strand DNA nucleotides in the full-transcription-bubble nucleic-acid scaffold (Figures 1C-D and 3B).

### Cryo-EM structure of RPrtc,≥11: RNA exit through nontemplate-strand scrunching portal

In the cryo-EM structure of RPrtc,≥11 [C_+1_C_+2_C_+3_], the RNA segment 6-11 nt from the RNA 3’ end (nucleotides rN6-rN11) follows a path that differs by ∼130° from the path of the RNA product in standard transcription initiation and elongation complexes, and that differs by ∼30° from the paths of the shorter RNA products in the crystal structures of RPrtc in Murakami et al., 2017 and Shin et al., 2020 (Figures 3A-B,D and 4C). This RNA segment, nucleotides rN6-rN11, diverges from the path of the RNA product in standard transcription initiation and elongation complexes, due to a collision with the σ-finger tip, involving nucleotide rN6 (Figure 3B,D and 4C, left). This RNA segment, nucleotides rN6-rN11, then cross the transcription bubble, spanning ∼20 Å, cross the DNA nontemplate strand, spanning an additional ∼10 Å, and exit from the RNAP active-center cleft at a position on the face of RNAP opposite the RNA exit used in standard transcription elongation complexes (Figures 3B,D and 4C, center and right).

Three residues of the σ-finger tip make van der Waals interactions with nucleotide rN6, causing the path of nucleotides rN6-rN11 to diverge, by ∼130°, from the path of the RNA product in a standard transcription initiation or elongation complex (σ residues D323, E324, and D326; residues numbered here and below as in *T. thermophilus* RNAP holoenzyme; Figure 4C, left). Four residues of RNAP β subunit and one residue of σ make H-bonded or salt-bridged interactions with the sugar-phosphate backbone of nucleotides rN6-rN11 (β residues K188, R189, T419 and R420, and σ residue K325; Figure 4C). Two residues of RNAP β subunit and one residue of σ make single H-bonds with RNA bases of nucleotides rN6-rN11 (β residues N187 and G417, and σ residue T77; Figure 4C). The observations that most protein-RNA interactions with nucleotides rN6-rN11 involve the sugar-phosphate backbone, and that interactions with bases involve single H-bonds that, with wobble, could be made with any base, suggest that the RNA-exit pathway observed in this structure may be compatible with any RNA sequence. Residues of RNAP β subunit that make protein-RNA interactions with nucleotides rN6-rN11 are residues located immediately N-terminal to β conserved region βa5 (β residues 187-189) and residues located in β conserved region βa7, also known as fork-loop 2 (β residues 417-420) (Figure S3, left; β conserved regions defined as in Lane and Darst, 2010). All six residues of RNAP β subunit that interact with nucleotides rN6-rN11 are invariant or highly conserved across Gram-negative, Gram-positive, and *Thermus*-*Deinococcus*-clade bacteria (Figure S3, left). Residues of σ that make protein-RNA interactions with nucleotides rN6-rN11 are residues at the N-terminus of σR1.2 (σ residue 77) and residues of the part of σR3.2 that forms the σ-finger tip (σ residues 323-326) (Figure S3, right; β conserved regions defined as in Feklistov et al., 2014). Most, three of five, residues of σ that interact with nucleotides rN6-rN11 are invariant or highly conserved across Gram-negative, Gram-positive, and *Thermus*-*Deinococcus*-clade bacteria (Figure S3B). The observation that residues of RNAP β and σ that make protein-RNA interactions with nucleotides rN6-rN11 are highly conserved across bacterial species suggest that the RNA-exit pathway observed in this structure may mediate the production of long RNA products by reiterative transcription initiation across bacterial species.

At the point where nucleotides rN6-rN11 cross the DNA nontemplate strand, direct DNA-RNA interactions occur, involving the stacking of the base of the nucleotide at nontemplate-strand position +2 (8 nt downstream of −10 element) on the base of nucleotide rN8 (Figure 3B,D and 4C, center). This DNA-RNA base-stacking interaction is facilitated by the 1 nt of scrunching of the nontemplate strand, which, by unstacking and flipping out the nucleotide at nontemplate-strand position +1, frees the nucleotide at nontemplate-strand position +2 for DNA-RNA base stacking (Figure 3B,D and 4C, center). The ability of nucleotides rN6-rN11 to cross the DNA nontemplate strand is further facilitated by the 1 nt of scrunching of the nontemplate strand in that the unstacking and flipping of the nucleotide at nontemplate strand position +1 opens space for, and removes a steric barrier to. passage of nucleotides rN8 and rN9 past the DNA nontemplate strand.

Nucleotides rN8-rN10 are accommodated within the same cavity within the RNAP active-center cleft that accommodates scrunched nontemplate-strand DNA in standard transcription initiation (Figure 3A-B,D; Basu et al., 2014; Hasemeyer, 2018; Kapanidis et al., 2006; Winkelman et al., 2015). Nucleotide rN11 exits the RNAP active-center cleft and interacts with bulk solvent. Nucleotide rN11 exits the RNAP active-center cleft through the same opening through which long, ≥6-8 nt, segments of scrunched nontemplate-strand DNA exit the RNAP active-center cleft (alternative RNA exit in Figure 3A-B; nontemplate-strand scrunching portal in Hasemeyer, 2018; Kapanidis et al., 2006; Winkelman et al., 2015). We infer that production of long RNAs by reiterative transcription initiation exploits the same cavity within the RNAP active-center cleft (providing an alternative RNA path) and the same opening from the RNAP active-center cleft (providing an alternative RNA exit) that exist to accommodate and extrude scrunched nontemplate-strand DNA in standard transcription initiation.

Two changes in RNAP conformation present in the structure obtained following reiterative transcription initiation in solution (Figures 3-4), but not in structures obtained following reiterative transcription initiation *in crystallo* (Figures 1-2; Murakami et al., 2017; Shin et al., 2020), account for the ability to produce long RNAs in the former, but not in the latter. First, in the structure of RPrtc,≥11 [C_+1_C_+2_C_+3_], the tip of the σ finger folds back on itself, moving ∼3 Å farther away from the RNAP active center (Figures 3B, right and 4D), in a manner similar to, but less marked than, the folding back of the tip of the σ finger, driven by collision with the RNA 5’ end, that occurs in standard transcription initiation upon extension of the RNA product to a length of 6 nt (Figure 4D; Li et al., 2020). This change in local conformation of the σ-finger tip enables reiteratively transcribed RNA to enter into the alternative RNA pathway and to be extended beyond a length of 5 nt. Second, in the structure of RPrtc,≥11 [C_+1_C_+2_C_+3_], the RNAP clamp (Chakraborty et al., 2012; Duchi et al., 2018; Mazumder et al., 2020) opens by ∼3°, increasing the width of the RNAP active-center cleft by ∼2 Å (Figure 4E). This change in RNAP clamp conformation enables reiteratively transcribed RNA to cross nontemplate-strand DNA in order to access the alternative RNA exit and leave the RNAP active-center cleft. In our crystal structures of complexes obtained following reiterative transcription initiation *in crystallo*, neither of these two conformational changes occurred, and RNA products therefore were limited to lengths of 4-5 nt (Figure 1C-D). In other crystal structures obtained following reiterative transcription initiation *in crystallo*, using different sequences and different conditions, the first, but not the second, of these two conformational changes occurred, and RNA products therefore were limited to lengths of 6-8 nt (Murakami et al., 2017; Shin et al., 2020).

### Mapping of RNAP leading-edge and trailing-edge positions in RPrtc: RNA extension without DNA scrunching

DNA scrunching by RNAP has two biochemically detectable hallmarks: (i) downstream movement of the RNAP leading edge, but not the RNAP trailing edge, relative to DNA (Kapanidis et al., 2006; Winkelman et al., 2016a; Winkelman and Gourse, 2017; Winkelman et al., 2016b; Yu et al., 2017), and (ii) expansion of the transcription bubble (Revyakin et al., 2006; Yu et al., 2017). In a preceding section, we proposed that RNA extension in reiterative transcription does not involve DNA scrunching, except for 1 bp of DNA scrunching to position the 3’ end of the initial RNA product in the RNAP active-center P site. As a first approach to test this proposal, we assessed positions of the RNAP leading edge and trailing edge relative to DNA during reiterative transcription initiation (Figure 5).

**Figure 5.**
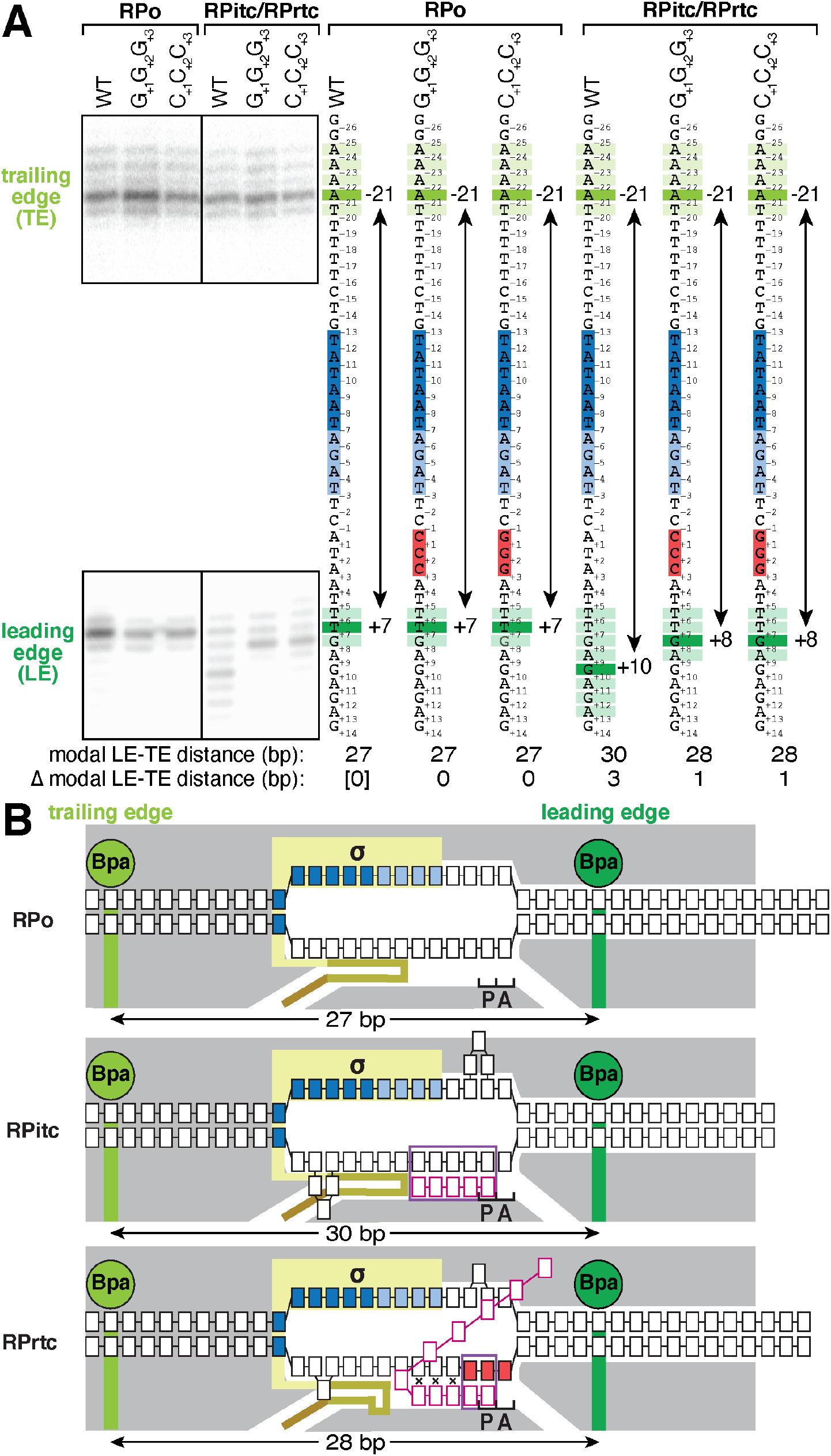
Mapping of RNAP leading-edge and trailing-edge positions in RPrtc by use of protein-DNA photocrosslinking: RNA extension without DNA scrunching. **(A)** RNAP trailing-edge and leading-edge positions in transcription initiation complexes at the N25 promoter (WT) and derivatives of the N25 promoter containing template-strand G_+1_G_+2_G_+3_ and C_+1_C_+2_C_+3_ homopolymeric sequences (G_+1_G_+2_G_+3_ and C_+1_C_+2_C_+3_). First bracketed subpanel, protein-DNA photocrosslinking data for RNAP-promoter open complex (RPo); second bracketed subpanel, protein-DNA photocrosslinking data for transcription initiation complexes engaged in standard transcription initiation (RPitc) and reiterative transcription initiation (RPrtc); third bracketed subpanel, interpretation of protein-DNA photocrosslinking data for RPo; fourth bracketed subpanel, interpretation of protein-DNA photocrosslinking data for RPitc and RPrtc. In third and fourth subpanels, promoter sequences are shown with positions numbered relative to the transcription start site, and with positions of the −10-element, the discriminator element, and the homopolymeric sequence highlighted in blue, light blue, and red. Thick and thin light green bars indicate strong and weak RNAP-trailing-edge crosslinks, and thick and thin dark green bars indicate strong and weak RNAP-leading-edge crosslinks. Bottom of panel, observed modal trailing-edge/leading-edge distances (modal TE-LE distance) and differences in modal TE-LE distance relative to modal TE-LE distance in RPo at wild-type N25 promoter [Δ(modal TE-LE distance)]. **(B)** Mechanistic interpretation of data in panel A. Three states are shown: RPo, RPitc [specifically, RPitc having a 5 nt RNA product in a post-translocated state (RPitc,5 post), corresponding to the major crosslink in panel A], and RPrtc. Gray, RNAP; yellow, σ; yellow-brown, σ finger (note displacement of σ-finger tip in RPrtc); brown, σ region-3/region-4 linker; light green, trailing-edge Bpa and crosslinking site for trailing-edge Bpa; dark green, leading-edge Bpa and crosslinking site for leading-edge Bpa; black boxes with blue fill, −10-element nucleotides; black boxes with light blue fill, discriminator-element nucleotides; black boxes with red fill, template-strand homopolymeric-sequence nucleotides; other black boxes, other DNA nucleotides (nontemplate-strand nucleotides above template-strand nucleotides); magenta boxes, RNA nucleotides; violet rectangles, RNA-DNA hybrids; P and A, RNAP active-center product and addition sites. Raised template-strand nucleotides and black x’s indicate non-base-paired nucleotides. Scrunching of nontemplate and template DNA strands is indicated by bulged-out nucleotides. Initial-product formation in both standard transcription initiation and reiterative transcription initiation involves one step of DNA scrunching. RNA extension in standard transcription initiation involves additional DNA scrunching, but RNA extension in reiterative transcription does not.

We used unnatural amino-acid mutagenesis to incorporate the photoactivatable amino acid p-benzoyl-L-phenylalanine (Bpa) at the RNAP leading edge and trailing edge, and we used protein-DNA photocrosslinking to define positions of the RNAP leading edge and trailing edge relative to DNA (methods essentially as in Winkelman et al., 2016b; Yu et al., 2017). We analyzed reiterative transcription initiation at derivatives of the bacteriophage N25 promoter containing either a template-strand G_+1_G_+2_G_+3_ homopolymeric sequence (RPo [G_+1_G_+2_G_+3_] and RPrtc [G_+1_G_+2_G_+3_]) or a template-strand C_+1_C_+2_C_+3_ homopolymeric sequence (RPo [C_+1_C_+2_C_+3_] and RPrtc [C_+1_C_+2_C_+3_]). For reference, we analyzed standard transcription initiation at the wild-type N25 promoter (RPo WT and RPitc WT). *In vitro* transcription experiments, carried out using the same reaction conditions, show production of long, up to at least 50 nt, reiterative transcripts with the G_+1_G_+2_G_+3_ and C_+1_C_+2_C_+3_ promoters in the presence of CTP and GTP, respectively, and show production of short, up to ∼8 nt, standard transcripts with the WT promoter in the presence of ATP and UTP (Figure S1B; Hsu et al., 2003; Revyakin et al., 2006).

The photocrosslinking results indicate that RPrtc exhibits an RNAP trailing-edge position that is unchanged as compared to RPo, exhibits an RNAP leading-edge position that is shifted downstream by 1 bp as compared to RPo of only 1 bp, and thus exhibits an RNAP trailing-edge/leading-edge distance that is increased by 1 bp as compared to RPo (Figure 5A). In contrast, RPitc exhibits an RNAP trailing-edge position that is unchanged as compared to RPo, exhibits an RNAP leading-edge position that is shifted downstream by up to 6 bp (range = 1-6 bp; mode = 3 bp) as compared to RPo, and thus exhibits an RNAP trailing-edge/leading-edge distance that is increased by up to 6 bp as compared to RPo (range of increase = 1-6 bp; modal increase = 3 bp; Figure 5A). An increase in RNAP trailing-edge/leading-edge distance is a defining hallmark of DNA scrunching (Kapanidis et al., 2006; Winkelman et al., 2016b; Yu et al., 2017). Thus, the results indicate that RPrtc engaged in synthesis of RNA products up to at least 50 nt in length exhibits only 1 bp of DNA scrunching, whereas RPitc engaged in synthesis of RNA up to 8 nt in length exhibits up to 6 bp of DNA scrunching. We conclude, consistent with the crystal structures of the preceding sections, that in reiterative transcription--following synthesis of a 2 nt initial RNA product and 1 bp of DNA scrunching to position the 3’ end of the initial RNA product in the RNAP active-center P site--RNA extension does not involve DNA scrunching (Figure 5B).

### Measurement of transcription bubble size in RPrtc: RNA extension without DNA scrunching

As a second approach to test the proposal that RNA extension in reiterative transcription does not involve DNA scrunching--except for 1 bp of DNA scrunching to position the 3’ end of the initial RNA product in the RNAP active-center P site--we assessed transcription-bubble expansion during reiterative transcription initiation (Figure 6).

**Figure 6.**
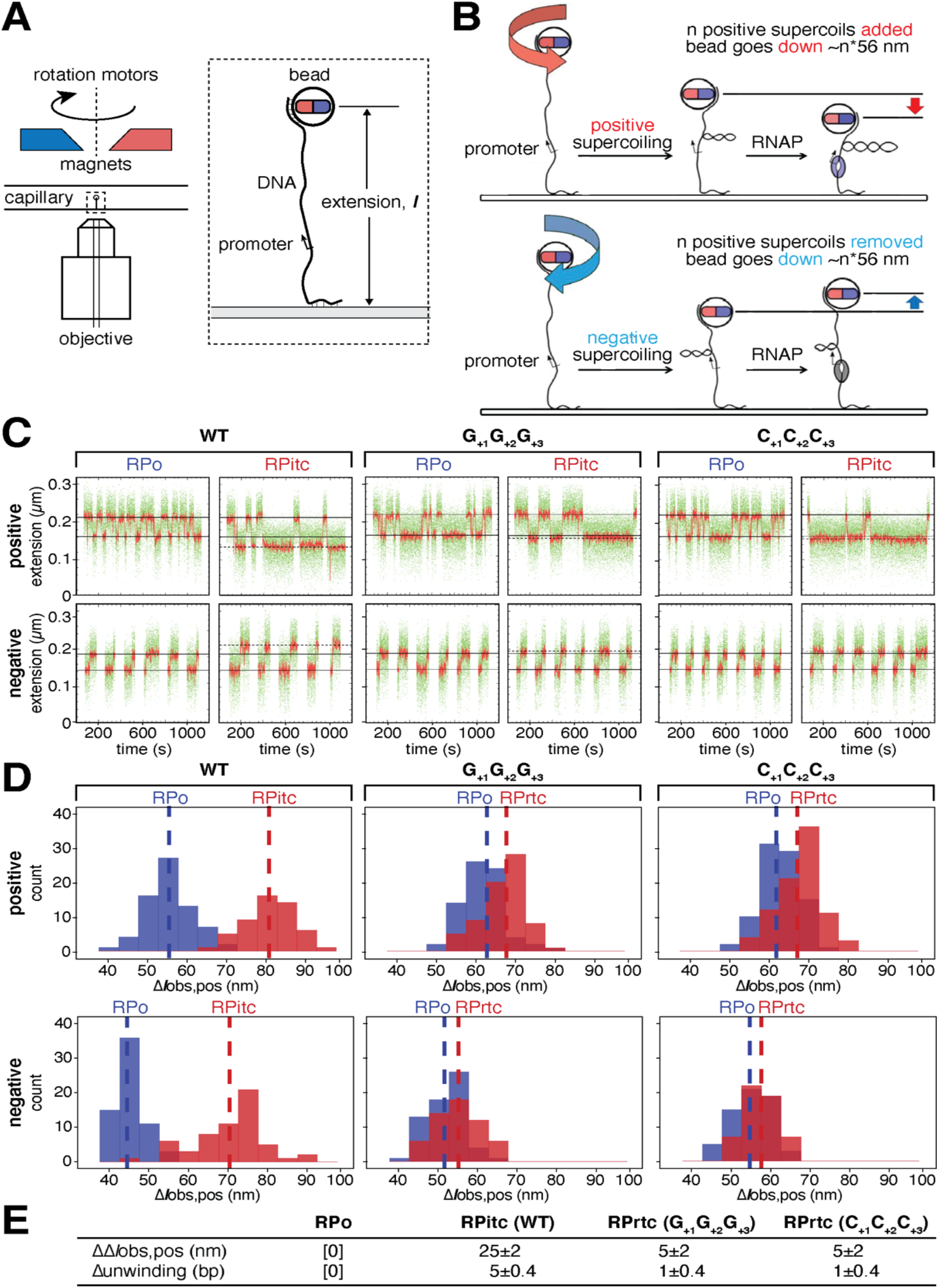
Measurement of transcription bubble size in RPrtc by use of single-molecule DNA nanomanipulation: RNA extension without DNA scrunching. **(A)-(B)** Experimental approach (Revyakin et al., 2005; Revyakin et al., 2006; Yu et al., 2017). (A), apparatus. (B), end-to-end extension (*l*) of a mechanically stretched, positively supercoiled (top), or negatively supercoiled (bottom), DNA molecule is monitored. Unwinding of n turns of DNA by RNAP results in compensatory gain of n positive supercoils or loss of n negative supercoils, and movement of the bead by n*56 nm. **(C)** Single-molecule time traces for RPo and RPitc at the N25 promoter (WT; left), and for RPo and RPrtc at derivatives of the N25 promoter containing template-strand G_+1_G_+2_G_+3_ and C_+1_C_+2_C_+3_ homopolymeric sequences (G_+1_G_+2_G_+3_ and C_+1_C_+2_C_+3_; middle and right). Upper subpanels, positively supercoiled DNA; lower subpanels, negatively supercoiled DNA. Green points, raw data (30 frames/s); red points, averaged data (1 s window); horizontal black lines, unbound and RPo states; dashed horizontal black lines, RPitc and RPrtc states (with the difference in Δ*l*obs between RPo and RPitc being substantially greater than the difference in Δ*l*_obs_ between RPo and RPrtc). **(D)** Single-molecule transition-amplitude histograms for RPo and RPitc at the N25 promoter (WT; left), and for RPo and RPrtc at derivatives of the N25 promoter containing template-strand G_+1_G_+2_G_+3_ and C_+1_C_+2_C_+3_ homopolymeric sequences (G_+1_G_+2_G_+3_ and C_+1_C_+2_C_+3_; middle and right). Upper subpanels, positively supercoiled DNA; lower subpanels, negatively supercoiled DNA. Vertical dashed lines, means; Δ*l*obs,pos, transition amplitudes with positively supercoiled DNA; Δ*l*obs,neg, transition amplitudes with negatively supercoiled DNA. **(E)** Differences in Δ*l*_obs,pos_ and DNA unwinding relative to those in RPo at wild-type N25 promoter (means ± 2SEM).

We used magnetic-tweezers single-molecule DNA-nanomanipulation to assess reiterative transcription (Figure 6A-B), analyzing the same promoter derivatives as in the preceding section (Figures 5, S1B). The resulting transition amplitudes, transition-amplitude histograms, and RNAP-dependent DNA unwinding values show that RNAP-dependent DNA unwinding is greater by 1±0.4 bp in RPrtc engaged in synthesis of RNA products up to at least 50 nt in length than in RPo (Figure 6C-E). In contrast, RNAP-dependent DNA unwinding is greater by 5±0.4 bp in RPitc engaged in synthesis of RNA products up to ∼8 nt than in RPo (Figure 6C-E). An increase in transcription-bubble size is a defining hallmark of DNA scrunching (Revyakin et al., 2006; Yu et al., 2017). Thus, the results from single-molecule DNA nanomanipulation indicate that RPrtc engaged in synthesis of RNA products up to at least 50 nt in length exhibits only 1 bp of DNA scrunching. We conclude, consistent with the conclusions of the preceding sections, that, in reiterative transcription--following synthesis of a 2-nt initial RNA product and 1 bp of DNA scrunching to position the 3’ end of the initial RNA product in the RNAP active-center P site--RNA extension does not involve DNA scrunching.

## Discussion

### Mechanism of reiterative transcription

Taken together, our results from x-ray crystallography (Figures 1-2), cryo-EM (Figures 3-4), site-specific protein-DNA photocrosslinking (Figure 5), and DNA single-molecule nanomanipulation (Figure 6) establish that, whereas standard transcription initiation involves 1 bp of DNA scrunching for each step of RNA extension, reiterative transcription initiation at promoters containing template-strand G_+1_G_+2_G_+3_ and C_+1_C_+2_C_+3_ homopolymeric sequences involves only 1 bp of DNA scrunching, irrespective of the number of steps of RNA extension. We conclude that, in reiterative transcription initiation at promoters containing template-strand G_+1_G_+2_G_+3_ and C_+1_C_+2_C_+3_ homopolymeric sequences, following synthesis of the initial 2 nt RNA product (Figure 7, left, lines 1-2), 1 bp of DNA scrunching occurs to place the RNA 3’ end in the RNAP active-center P site (Figure 7, left, lines 2-3), and no additional DNA scrunching occurs in RNA extension (Figure 7, right). We infer that, in reiterative transcription initiation, RNA extension does not involve movement of DNA relative to the RNAP active center, but, instead, involves RNA slipping, in which, in each step of RNA extension, RNA slips upstream by 1 nt relative to both template-strand DNA and the RNAP active center to place the RNA 3’ end in the RNAP active-center P site (Figure 7, right). Thus, we infer that, in RNA extension in reiterative transcription initiation, the nucleotide addition cycle consists of RNA slipping to convert a pre-translocated state having a 3 bp RNA-DNA hybrid and having the RNA 3’ end in the RNAP active-center A site (“pre-slipped” state; Figure 7, left, RPitc,3 pre and Figure 7, right, RPrtc,n pre) into a post-translocated state having a 2 bp RNA-DNA hybrid and having the RNA 3’ end in the RNAP active-center P site (“post-slipped” state; Figure 7, right, RPrtc,n post), followed by NTP binding, phosphodiester bond formation, and pyrophosphate release (Figure 7, right). Thus, in contrast to standard transcription initiation, in which the RNA-DNA hybrid increases in length by 1 bp in each step of RNA extension, up to a length of ∼10 bp, at which point promoter escape ensues (Figure 7, left; Hsu et al., 2003; Mazumder and Kapanidis, 2019; Ruff et al., 2015; Winkelman et al., 2021), in reiterative transcription initiation, the RNA-DNA hybrid does not extend beyond a 3 bp state, and, instead, alternates between a 3 bp pre-slipped state and a 2 bp post-slipped state in each step of RNA extension (Figure 7, right). RNA slipping by 1 nt breaks 1 bp of the RNA-DNA hybrid--specifically the upstream-most base pair of the RNA-DNA hybrid (Figure 7, right); breakage of the upstream-most base pair of the RNA-DNA hybrid occurs because slipping moves the RNA nucleotide that had been base-paired to the upstream-most nucleotide of the template-strand homopolymeric sequence into alignment with a non-complementary DNA nucleotide upstream of the template-strand homopolymeric sequence (Figure 7, right). According to this mechanism, the “branching point,” or “decision point,” between the standard-transcription-initiation and reiterative-transcription-initiation pathways occurs upon formation of RPitc,3 pre (Figure 7, lines 4-5; at this decision point, DNA scrunching followed by RNA extension yields standard transcription initiation, whereas RNA slipping followed by RNA extension yields reiterative transcription initiation (Figure 7, lines 4-5).

**Figure 7.**
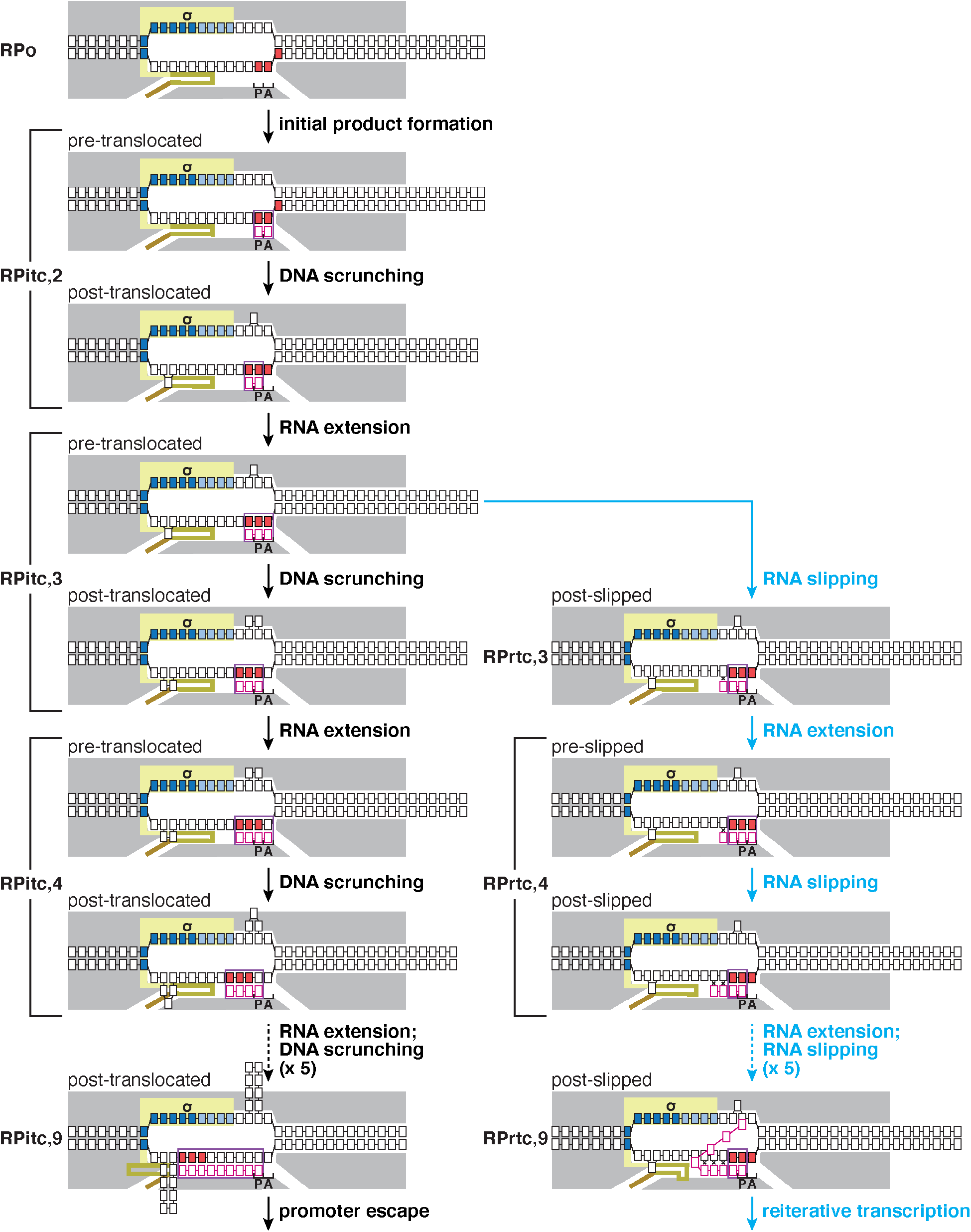
Mechanisms of standard transcription initiation and reiterative transcription initiation. Standard transcription initiation (left column) and reiterative transcription initiation (first four panels of left column followed by panels of right column). Cyan, reactions present only in reiterative transcription: i.e., cycles of RNA extension and slippage. Other colors and symbols are as in Figure 5. Scrunching is indicated by bulged-out nucleotides (∼8-10 scrunched bp prior to promoter escape in the standard transcription initiation pathway; 1 scrunched bp in the reiterative transcription initiation pathway). Scrunched nucleotides of nontemplate and template DNA strands during initial transcription are accommodated as bulges within the unwound transcription bubble.

The mechanism for reiterative trascription initiation defined here and set forth in Figure 7, requires a template-strand homopolymeric sequence at least 3 nt in length. This aspect of the mechanism is consistent with, and supported by, the observation from previous work that reiterative transcription initiation is efficient only at promoters containing template-strand homopolymeric sequences at least 3 nt in length (Cheng et al., 2001; Turnbough and Switzer, 2008; Turnbough Jr, 2011; Vvedenskaya et al., 2015; Xiong and Reznikoff, 1993).

Our results from cryo-EM structure determination of RPrtc,≥11 [C_+1_C_+2_C_+3_] (Figures 3-4) further show that production of long, ≥11 nt, RNAs by reiterative transcription initiation involves an alternative RNA path for nucleotides rN6-rN11 and an alternative RNA exit for nucleotides rN≥11 (Figures 3A-B,D and 4C). The alternative RNA path exploits the cavity within the RNAP active-center cleft that accommodates scrunched nontemplate-strand DNA in standard transcription initiation (Figure 1B; Basu et al., 2014; Hasemeyer, 2018; Kapanidis et al., 2006; Winkelman et al., 2015), and the alternative RNA exit exploits the portal between the RNAP active-center cleft and bulk solvent that mediates the extrusion of long segments of scrunched nontemplate-strand DNA in standard transcription initiation (Hasemeyer, 2018; Kapanidis et al., 2006; Winkelman et al., 2015). Because reiterative transcription initiation involves only 1 bp of DNA scrunching (Figures 1C-D, 3A-C, and 5-7), this cavity within the RNAP active-center cleft and this portal between the RNAP active-center cleft and bulk solvent--both of which would be occupied by scrunched nontemplate-strand DNA in standard transcription initiation--are not occupied by scrunched nontemplate-strand DNA in reiterative transcription and, instead, are available to be occupied by RNA.

A small change in local conformation of the σ-finger tip--a folding back of the σ-finger tip upon itself--enables reiteratively transcribed RNA to enter the alternative RNA path, thereby enabling the extension of reiteratively transcribed RNA beyond a length of 5 nt (Figure 4D). A small change in RNAP clamp conformation--an opening of the RNAP clamp by ∼3°--enables reiteratively transcribed RNA to cross the nontemplate DNA strand and to access the alternative RNA exit, thereby enabling the extension of reiteratively transcribed RNA beyond a length of 8 nt and the extrusion of reiteratively transcribed RNA from the RNAP active-center cleft (Figure 4E).

Because the alternative RNA path and the alternative RNA exit enable reiteratively transcribed RNA to be extruded from the RNAP active-center cleft without substantially disrupting σ-DNA and σ-RNAP interactions (Figure 3A-B), these features enable reiterative transcription initiation to generate long RNAs without promoter escape.

Because most protein-RNA interactions in the alternative RNA path and alternative RNA exit involve interactions with the RNA sugar-phosphate backbone (Figure 4C), and because interactions with RNA bases involve only single H-bonds that likely could be made, with wobble, by any RNA nucleotide (Figure 4C), the alternative RNA path and alternative RNA exit are likely to be compatible with any RNA sequence.

Because the RNAP and σ residues that make protein-RNA interactions with RNA in the alternative RNA path and alternative RNA exit are invariant or highly conserved across Gram-negative, Gram-positive, and *Thermnus*-*Deinococcus*-clade bacteria (Figure S3), the alternative RNA path and alternative RNA exit, and the mechanism of reiterative transcription initiation set forth here and in Figure 7, are likely to be conserved across bacterial species.

### Prospect

The results and mechanism of this work, pertain to reiterative transcription initiation at promoters containing template-strand G_+1_G_+2_G_+3_ and C_+1_C_+2_C_+3_ homopolymeric sequences (Figures 1-7). Previous work shows that reiterative transcription initiation also occurs efficiently at promoters containing template-strand A_+1_A_+2_A_+3_ and T_+1_T_+2_T_+3_ homopolymeric sequences (Turnbough and Switzer, 2008; Turnbough Jr, 2011; Vvedenskaya et al., 2015) and at promoters containing ≥ 4 nt homopolymeric sequences (Turnbough and Switzer, 2008; Turnbough Jr, 2011; Vvedenskaya et al., 2015). We consider it likely that the mechanism of Figure 7 also applies to these promoters. Previous work also shows that reiterative transcription initiation can occur at promoters containing more complex, non-homopolymeric repeat sequences (Turnbough and Switzer, 2008; Turnbough Jr, 2011; Vvedenskaya et al., 2015). We consider it likely that a mechanism related to the mechanism of Figure 7, but with different extents of DNA scrunching and RNA slipping, applies at these promoters. We note that the crystal-structure determination, cryo-EM structure determination, site-specific protein-DNA photocrosslinking, and single-molecule nanomanipulation procedures of this report could be used to define mechanisms of reiterative transcription at any promoter.

Previous work has shown that complexes engaged in reiterative transcription initiation synthesizing RNA products up to at least 8 nt in length can switch from reiterative transcription initiation to standard transcription initiation, yielding productive complexes that escape the promoter and synthesize full-length RNA products comprising non-templated, reiterative-transcription-dependent nucleotides at their 5’ ends followed by templated, standard-transcription-dependent nucleotides (Turnbough and Switzer, 2008; Turnbough Jr, 2011; Vvedenskaya et al., 2015). Key unresolved questions include how this switching occurs and where the RNA products that result from switching exit the RNAP active-center cleft. We hypothesize that RNA products that result from switching exit the RNAP active-center cleft through the RNAP RNA-exit channel, with the templated RNA segment generated after switching to standard transcription initiation proceeding into, and through, the RNAP RNA-exit channel as in standard transcription, and pulling the 5’ non-templated RNA segment behind it. According to this hypothesis, the RNA product that results from switching would proceed into, and through, the RNAP RNA-exit channel as an RNA loop, and would trigger promoter escape by displacing the σ region-3/region-4 linker from the RNA-exit channel as in standard transcription (Figure S4, right). Cryo-EM structures of transcription complexes containing double-stranded RNA in the RNAP RNA-exit channel verify that the dimensions of the RNAP RNA-exit channel could accommodate such an RNA loop (Guo et al., 2018; Kang et al., 2018). Cryo-EM structure determination, analyzing transcription elongation complexes containing 5’ non-templated RNA segments generated by switching from reiterative transcription initiation to standard transcription initiation, potentially could provide a means to test this hypothesis.

## Acknowledgements

Work was supported by National Institutes of Health grants GM118059 (BEN) and GM041376 (RHE) and National Natural Science Foundation of China grant 31822001 (YZ).

## Author Contributions

Conceptualization, Y.L., J.T.W., L.Y., Y.Z., B.E.N., and R.H.E.; Investigation, Y.L., J.T.W., L.Y., C.P., E.F., J.T.K., and Y.Z.; Formal Analysis, Y.L., J.T.W., L.Y., and Y.Z.; Writing, Y.L., J.T.W., B.E.N., and R.H.E.; Visualization, Y.L., J.T.W., B.E.N., and R.H.E.; Supervision, B.E.N. and R.H.E.; Project Administration, B.E.N. and R.H.E.; Funding Acquisition, Y.Z., B.E.N. and R.H.E.

## Supplementary Materials

### Materials and Methods

#### Proteins

*T. thermophilus* RNAP core enzyme was prepared from *T. thermophilus strain* H8 (DSM 579; German Collection of Microorganisms and Cell Cultures GmbH), using cell lysis, polyethylenimine precipitation, ammonium sulfate precipitation, and ion-exchange chromatography on SP Sepharose FF (GE Healthcare Life Sciences, Inc.), Mono Q (GE Healthcare Life Sciences, Inc.), and Mono S HR (GE Healthcare Life Sciences, Inc.), as in (Maffioli et al., 2017; Zhang et al., 2012).

*T. thermophilus* σ^A^ was prepared by expression of a gene for N-terminally hexahistidine-tagged *T. thermophilus* σ^A^ in *E. coli*, followed by cell lysis, immobilized-metal-ion affinity chromatography on Ni-NTA agarose (Qiagen, Inc.), and anion-exchange chromatography on Mono Q (GE Healthcare Life Sciences, Inc.), as in (Zhang et al., 2012).

*T. thermophilus* RNAP σ^A^ holoenzyme was prepared by incubating 15 μM *T. thermophilus* RNAP core enzyme and 60 μM *T. thermophilus* σ^A^ in 2 ml 20 mM Tris-HCl (pH 7.9), 150 mM NaCl, and 1% glycerol for 12 h at 4°C, followed by size-exclusion chromatography on Superdex 200 (GE Healthcare Life Sciences, Inc.), as in (Zhang et al., 2012).

Bpa-containing *E. coli* RNAP core enzyme derivatives RNAP**-**β’ ^R1148Bpa^ and RNAP**-**β’ ^T48Bpa^ were prepared from *E. coli* strain NiCo21(DE3) (New England Biolabs, Inc.) transformed with plasmid pEVOL-pBpF (Chin et al., 2002) and either plasmid pIA900-RNAP**-**β’ ^R1148Bpa^ (Winkelman et al., 2015) or plasmid pIA900-RNAP**-**β’ ^T48Bpa^ (Winkelman et al., 2015), using procedures as in (Winkelman et al., 2015). *E. coli* RNAP core enzyme without Bpa was prepared from *E. coli* strain NiCo21(DE3) (New England Biolabs, Inc.) transformed with plasmid pIA900 (Svetlov and Artsimovitch, 2015), using procedures as in (Winkelman et al., 2015).

*E. coli* σ^70^ was prepared from *E. coli* strain NiCo21(DE3) transformed with plasmid pσ^70^-His (gift of J. Roberts; Marr and Roberts, 1997), using procedures as in (Marr and Roberts, 1997).

*E. coli* RNAP σ^70^ holoenzyme was prepared by incubating 1 μM *E. coli* RNAP core enzyme and 5 μM *E. coli* σ^70^ in 10 mM Tris-Cl (pH 8.0), 100 mM KCl, 10 mM MgCl_2_, 0.1 mM EDTA, 1 mM DTT, and 50% glycerol for 30 min at 25°C.

#### Oligodeoxyribonucleotides

Oligodeoxyribonucleotides (Table S2) were purchased from IDT, Inc.

#### Nucleic-acid scaffolds for structure determination

Nucleic-acid scaffolds for crystallization were prepared by incubating 20 nmol nontemplate-strand oligodeoxyribonucleotide and 22 nmol template-strand oligodeoxyribonucleotide in 50 μl 5 mM Tris-HCl (pH 7.9), 200 mM NaCl, and 10 mM MgCl_2_ for 5 min at 95°C, followed by cooling to 25°C in 2°C steps with 1 min per step using a thermal cycler (Applied Biosystems, Inc.), and were stored at −80°C.

#### Crystal-structure determination: crystal growth and in crystallo RNA synthesis

RPo was prepared by mixing 20 μl 18 μM *T. thermophilus* RNAP σ^A^ holoenzyme in 20 mM Tris-HCl (pH 7.9), 100 mM NaCl, and 1% glycerol with 1.1 μl 0.4 mM nucleic-acid scaffold in 5 mM Tris-HCl (pH 7.9), 200 mM NaCl, and 10 mM MgCl_2_, and incubating 1 h at 25°C.

Crystallization was performed essentially as in (Zhang et al., 2012). Crystallization drops containing 1 μl 20 μM RPo in 20 mM Tris-HCl (pH 7.9), 100 mM NaCl, 1% glycerol and 1 μl reservoir solution [0.1 M Tris-HCl (pH 8.4), 180 mM KCl, 50 mM MgCl_2_, and 9% (m/v) PEG 4000] were equilibrated with 400 μl reservoir solution in sealed hanging-drop plates. Crystals were grown 5 days at 22°C. Micro-seeding was performed as needed using the same reservoir solution.

*In crystallo* RNA synthesis was performed by adding 0.2 μl 10 mM each NTP (ATP, UTP, and CTP for RPitc,5; CTP for RPrtc,5 [G_+1_G_+2_G_+3_]; GTP for RPrtc,4 [C_+1_C_+2_C_+3_]) in reservoir solution directly into crystallization drops at 22°C, incubating 2 min at 22°C, transferring crystals to 2 μl reservoir solution containing 18% (v/v) (2R, 3R)-(-)-2,3-butanediol (Sigma-Aldrich, Inc.) at 22°C, and flash-cooling in liquid nitrogen.

#### Crystal-structure determination: data collection and data reduction

Diffraction data were collected from cryo-cooled crystals at Stanford Synchrotron Radiation Lightsource (SSRL) beamline 9-2 (RPrtc,5 [G_+1_G_+2_G_+3_] and RPrtc,4 [C_+1_C_+2_C_+3_]; Table S2) and at Argonne Photon Source (APS) beamline 19-ID (RPitc,5; Table S2). Data were processed using HKL3000 (Minor et al., 2006). Structures were solved by molecular replacement using the crystal structure of *T. thermophilus* RPo (PDB 4G7H; Zhang et al., 2012) as search model. Iterative cycles of reciprocal refinement using Phenix (Liebschner et al., 2019) and real-space model building using Coot (Emsley et al., 2010) were performed. Improvement of the coordinate model resulted in improvement of phasing, and electron density maps for nucleic acids, which were not included in models at this stage of model building and refinement, improved over successive cycles. Nucleic acids then were built into the model and refined in stepwise fashion. The RPitc,5 model refined to R_work_ = 0.23 and R_free_ = 0.27, the RPrtc,5 [G_+1_G_+2_G_+3_] model refined to R_work_ = 0.24 and R_free_ = 0.29, and the RPrtc,4 [C_+1_C_+2_C_+3_] refined to R_work_ = 0.26 and R_free_ = 0.28. The final atomic models and structure factors were deposited in the Protein Data Bank (PDB) with accession codes RPitc,5 (7MLB), RPrtc,5 [G_+1_G_+2_G_+3_] (7MLI), and RPrtc,4 [C_+1_C_+2_C_+3_] (7MLJ; Table S2).

#### Cryo-EM structure determination: sample preparation

QUANTIFOIL R 2/1 Cu 300 mesh (Quantifoil) grids were glow-discharged and graphene-oxide-coated as in (Pantelic et al., 2010; Thomas et al., 2016). Grids were glow-discharged at 25 mA and 0.27 mBar for 60 s using a glow-discharge cleaning system (PELCO easiGlow; Ted Pella), 3 μl 0.2 mg/ml graphene oxide suspension in water (Sigma) was applied to each grid, and, after 90 s at 25°C, and were blotted using #595 filter paper (Ted Pella) for 1 s at 25°C, washed with 3×3 μl water (twice on graphene-oxide side and once on opposite side), and stored under vacuum at 25°C until use.

RPo was prepared by mixing 20 μl 18 μM *T. thermophilus* RNAP σ^A^ holoenzyme in 20 mM Tris HCl (pH 7.9), 100 mM NaCl, and 5 mM MgCl_2_ with 1.1 μl 0.4 mM nucleic-acid scaffold in 5 mM Tris HCl (pH 7.9), 200 mM NaCl, and 10 mM MgCl_2_ and incubating 1 h at 25°C. Samples of RPrtc were prepared by diluting 2 μl RPo with 16 μl 20 mM Tris HCl (pH 7.9), 100 mM NaCl, and 5 mM MgCl_2_ at 25°C, adding 2 ul 10 mM GTP, and incubating 20 min at 25°C. Aliquots of samples of RPrtc (3 μl) were applied to graphene-oxide-coated grids mounted on a Vitrobot Mark IV autoplunger (FEI/ThermoFisher) at 4°C and 95% relative humidity, and grids were incubated 30 s, blotted with #595 filter paper (Ted Pella) for 7 s, flash-frozen by plunging in liquid ethane cooled with liquid nitrogen, and stored in liquid nitrogen.

#### Cryo-EM structure determination: data collection and data reduction

Cryo-EM data were collected at the Rutgers University Cryo-EM and Nanoimaging Facility, using a 200 kV Talos Arctica (FEI/ThermoFisher) electron microscope equipped with a GIF Quantum K2 direct electron detector (Gatan). Data were collected using Serial EM (Mastronarde, 2018), a nominal magnification of 130,000x, a calibrated pixel size of 1.038 Å/pixel, and a dose rate of 4.97 electrons/pixel/s. Movies were recorded at 200 ms/frame for 6 s (30 frames). Defocus range was varied between −1.2 µm and −2.8 µm. A total of 3,424 micrographs were recorded from one grid over two days. Micrographs were gain-normalized and defect-corrected.

Data were processed as summarized in Figure S2A. Data processing was performed using a Tensor TS4 Linux GPU workstation with four GTX 1080 Ti graphic cards (NVIDIA). Dose-weighting motion correction (5×5 tiles; b-factor = 150) was performed using Motioncor2 (Zheng et al., 2017). Contrast-transfer-function (CTF) estimation was performed using CTFFIND-4.1 (Rohou and Grigorieff, 2015). Subsequent image processing was performed using Relion 3.1 (Zivanov et al., 2018). 2,143,284 Particles were picked using Relion Auto-picking with a 2D class reference. Particles were extracted into 256 pixel x 256 pixel boxes and were subjected to rounds of reference-free 2D classification, 3D classification, and removal of poorly populated classes, yielding a selected set of 663,667 particles. The selected set was used in 3D auto-refinement, CTF refinement (addressing higher-order aberrations, anisotropic magnification, and per-particle defocus), Bayesian polishing, and post-processing. A total of 339,426 particles was selected after further classification, and a density map at 3.0 Å overall resolution was obtained, as determined from gold-standard Fourier shell correlation (FSC; Figure S2D-E). The initial atomic model was built by manual rigid-body fitting of RNAP β’, RNAP β, RNAP α^I^, RNAP α^II^, RNAP ω, and σ segments from the crystal structure of *T. thermophilus* RPo (PDB 4G7H; Zhang et al., 2012) into the cryo-EM density map using Coot (Emsley et al., 2010), followed by manual fitting of DNA and RNA into the cryo-EM density map using Coot. For RNAP β’ residues 219-337 (part of the T. thermophilus β’ non-conserved region; Chlenov et al., 2005), density was absent, suggesting high segmental flexibility; this segment was not fitted. The complex was real-space-refined using secondary structure and nucleic acid restraints in PHENIX (Liebschner et al., 2019). The final atomic model, with map-to-model correlation of 0.70, was deposited in the Electron Microscopy Data Bank (EMDB) and the PDB with accession codes EMD-24424 and 7RDQ, respectively.

#### DNA templates for in vitro transcription and crosslinking assays

Linear DNA templates used for *in vitro* transcription (Figure S1B) and *in vitro* photo-crosslinking assays (Figure 5) contain sequences from positions −72 to +71 of N25 or positions −72 to +71 of N25 promoter containing template-strand G_+1_G_+2_G_+3_ and C_+1_C_+2_C_+3_ homopolymeric sequences (N25 [G_+1_G_+2_G_+3_] and N25 [C_+1_C_+2_C_+3_], respectively). Templates were generated by PCR in reactions containing 1 X Phusion HF Master Mix (Thermo Fisher Scientific, Inc.), 0.8 µM primer N25B, 0.8 µM primer N25T, and ∼1 pg of oligos JW173 (N25), JW169 (N25 [G_+1_G_+2_G_+3_]), or JW170 (N25 [C_+1_C_+2_C_+3_]). Reaction products were purified using a PCR purification kit (Qiagen, Inc.).

Nontemplate-strand sequence from positions −72 to +71 of N25:

5’-GAGAGAGGTACCTCGAGGGAAATCATAAAAAATTTATTTGCTTTCAGGAAAATTTTTCTGTATAATAGATTCATAATTTGAGAGAGGAGTTTAAATATGGCTGGTTCTCGCGAGAATTCCGAATAGCCATCCCAATCGAACAG-3’

Nontemplate-strand sequence from positions −72 to +71 of N25 [G_+1_G_+2_G_+3_]:

5’-GAGAGAGGTACCTCGAGGGAAATCATAAAAAATTTATTTGCTTTCAGGAAAATTTTTCTGTATAATAGATTCCCCATTTGAGAGAGGAGTTTAAATATGGCTGGTTCTCGCGAGAATTCCGAATAGCCATCCCAATCGAACAG-3’

Nontemplate-strand sequence from positions −72 to +71 of N25 [C_+1_C_+2_C_+3_]:

5’-GAGAGAGGTACCTCGAGGGAAATCATAAAAAATTTATTTGCTTTCAGGAAAATTTTTCTGTATAATAGATTCGGGATTTGAGAGAGGAGTTTAAATATGGCTGGTTCTCGCGAGAATTCCGAATAGCCATCCCAATCGAACAG-3’

#### In vitro transcription

*In vitro* transcription experiments in Figure S1A were performed using reaction mixtures (40 μl) containing 17 μM *T. thermophilus* RNAP σ^A^ holoenzyme or *E. coli* RNAP σ^70^ holoenzyme and 21 μM nucleic-acid scaffold (see “Nucleic-acid scaffolds for structure determination”) in 20 mM Tris-HCl (pH 7.9), 100 mM NaCl, and 5 mM MgCl_2_. For reactions with scaffold 1 (sequence in Figure S1A), 1 mM ATP and 1 mM UTP, along with 1 µl 200 Bq/fmol [ɑ^32^P]-UTP (PerkinElmer, Inc.), were added to initiate RNA synthesis. For reactions with scaffold 2 (sequence in Figure S1A), 1 mM CTP, along with 1µl 200 Bq/fmol [ɑ^32^P]-CTP (PerkinElmer, Inc.), was added to initiate RNA synthesis. For reactions with scaffold 3 and scaffold 4 (sequences in Figure S1A), 1 mM GTP, along with 1µl 200 Bq/fmol [ɑ^32^P]-GTP (PerkinElmer, Inc.), was added to initiate RNA synthesis. After 3 min at 25°C (for *T. thermophilus* RNAP σ^A^ holoenzyme) or 0.5 min at 25°C (for *E. coli* RNAP σ^70^ holoenzyme), reactions were terminated by addition of 100 µl 10 mM EDTA (pH 8.0) and 1 mg/ml glycogen.

*In vitro* transcription experiments in Figure S1B were performed using reaction mixtures (10 μl) containing 40 nM RNAP holoenzyme derivative and 4 nM promoter derivative (see “DNA templates for photocrosslinking assays”) in 10 mM Tris-HCl (pH 8.0), 100 mM KCl, 10 mM MgCl_2_, and 0.1 mg/ml bovine serum albumin. For reactions with promoter derivative N25 WT (sequence in Figure S1B), 200 µM ATP and 200 µM UTP, along with 1 µl 200 Bq/fmol [ɑ^32^P]-UTP (PerkinElmer, Inc.), were added to initiate RNA synthesis. For reactions with promoter derivative N25 [G_+1_G_+2_G_+3_] (sequence in Figure S1B), 200 µM CTP, along with 1µl 200 Bq/fmol [ɑ^32^P]-CTP (PerkinElmer, Inc.), was added to initiate RNA synthesis. For reactions with promoter derivative N25 [C_+1_C_+2_C_+3_] (sequence in Figure S1B), 200 µM GTP, along with 1µl 200 Bq/fmol [ɑ^32^P]-GTP (PerkinElmer, Inc.), was added to initiate RNA synthesis. After 20 min at 37°C, reactions were terminated by addition of 100 µl 10 mM EDTA (pH 8.0) and 1 mg/ml glycogen.

For both experiments in Figure S1A and experiments in Fiogire S1B, reaction mixtures were extracted with acidic phenol:chloroform (Ambion, Inc.), and RNA products were recovered by ethanol precipitation and re-suspended in 6.5 µl water. Samples were diluted 2-fold in urea loading dye (1 X TBE, 8 M urea, 0.025% xylene cyanol, and 0.025% bromophenol blue), and samples were analyzed by electrophoresis on 20%, 8 M urea, 1 X TBE polyacrylamide gels (UreaGel System; National Diagnostics, Inc.), followed by storage-phosphor imaging (Typhoon 9400 variable-mode imager; GE Healthcare Life Sciences, Inc.). Sizes of RNA products were estimated by comparison to radiolabeled Decade Marker (ThermoFisher Scientific, Inc.).

#### Determination of RNAP active-center A-site positions by protein-DNA photo-crosslinking in vitro

*In vitro* photo-crosslinking and crosslink mapping experiments were done using procedures described in (Yu et al., 2017). For the experiments in Figure 5, 50 μl reactions containing 100 nM RNAP holoenzyme, 10 nM template, 1 mM CpA dinucleotide (for N25 RPo samples), 1mM CpC dinucleotide (for N25 [G_+1_G_+2_G_+3_] RPo samples), or CpG dinucleotide (for N25 [C_+1_C_+2_C_+3_] RPo samples) and 200 µM ATP and UTP (for N25 RPitc samples), 200 µM CTP (for N25 [G_+1_G_+2_G_+3_] RPrtc samples), or 200 µM GTP (for N25 [C_+1_C_+2_C_+3_] RPrtc samples) and 1 X RB [10 mM Tris-Cl (pH 8.0), 70 mM NaCl, 10 mM MgCl_2_, and 0.1 mg/ml BSA] were incubated for 2 min at 25°C, and UV irradiated for 5 min at 25°C in a Rayonet RPR-100 photochemical reactor equipped with 16 x 350 nm tubes (Southern New England Ultraviolet, Inc.).

To denature RNAP-DNA complexes, reactions were mixed with 15 μl 5 M NaCl and 6 μl 100 µg/ml heparin, incubated for 5 min at 95°C, and cooled to 4°C. Crosslinked RNAP-DNA complexes were isolated by adding 20 µl MagneHis Ni-particles (Promega, Inc.) equilibrated and suspended in 10 mM Tris-Cl (pH 8.0), 1.2 M NaCl, 10 mM MgCl_2_, 10 μg/ml heparin, and 0.1 mg/ml BSA; MagneHis Ni-particles were collected using a magnetic microfuge tube rack; particles were washed with 50 µl 10 mM Tris-Cl (pH 8.0), 1.2 M NaCl, 10 mM MgCl_2_, 10 μg/ml heparin, and 0.1 mg/ml BSA, washed twice with 50 µl 1 X *Taq* DNA polymerase buffer (New England Biolabs, Inc.), and particles (which contained bound RNAP-DNA complexes) were resuspended in 10 µl 1 X *Taq* DNA polymerase buffer. For leading-edge crosslink mapping, primer extension reactions (12.5 µl) were performed by combining 2 µl of the recovered RNAP-DNA complexes, 1 µl of 1 µM ^32^P-5’-end-labeled primer N25B [200 Bq/fmol; prepared using [γ^32^P]-ATP (PerkinElmer, Inc.)] and T4 polynucleotide kinase (New England Biolabs, Inc.) as described in (Sambrook and Russell, 2006)], 1.25 μl 10 X dNTPs (2.5 mM dATP, 2.5 mM dCTP, 2.5 mM dGTP, 2.5 mM TTP; GE Healthcare Life Sciences, Inc.), 0.5 μl 5 U/μl Taq DNA polymerase (New England Biolabs, Inc.), 5 μl 5 M betaine, 0.625 μl 100% dimethyl sulfoxide, and 1.25 µl 10 X *Taq* DNA polymerase buffer; 40 cycles of 30 s at 95°C, 30 s at 53°C, and 30 s at 72°C. For trailing-edge crosslink mapping, primer extension reactions (12.5 µl) were performed by combining 2 µl of the recovered RNAP-DNA complexes, 1 µl of 1 µM ^32^P-5’-end-labeled primer N25T [200 Bq/fmol; prepared using [γ^32^P]-ATP (PerkinElmer, Inc.)] and T4 polynucleotide kinase (New England Biolabs, Inc.) as described in (Sambrook and Russell, 2006)], 1.25 μl 10 X dNTPs (2.5 mM dATP, 2.5 mM dCTP, 2.5 mM dGTP, 2.5 mM TTP; GE Healthcare Life Sciences, Inc.), 0.5 μl 5 U/μl Taq DNA polymerase (New England Biolabs, Inc.), 5 μl 5 M betaine, 0.625 μl 100% dimethyl sulfoxide, and 1.25 µl 10 X *Taq* DNA polymerase buffer; 40 cycles of 30 s at 95°C, 30 s at 53°C, and 30 s at 72°C. Reactions were stopped by addition of 12.5 μl 1 X TBE, 8 M urea, 0.025% xylene cyanol, and 0.025% bromophenol blue; radiolabeled products were separated by electrophoresis on 8%, 8 M urea, 1 X TBE polyacrylamide gels (UreaGel System; National Diagnostics, Inc.) and visualized by storage-phosphor imaging (Typhoon 9400 variable-mode imager; GE Healthcare Life Sciences, Inc.). Positions of RNAP-DNA crosslinking were determined by comparison to products of a DNA-nucleotide sequencing reaction generated using oligo N25T (when mapping trailing-edge crosslinks) or N25B (when mapping leading edge crosslinks) and a linear DNA template containing sequences from positions −72 to +71 of N25 (Thermo Sequenase Cycle Sequencing Kit; Affymetrix Inc.).

#### DNA templates for single-molecule DNA nanomanipulation

Plasmid pUC18-T20-N25-WT, which was used for experiments with the N25 promoter, was prepared as follows: The pUC18-T20 backbone was amplified by performing PCR using 1 ng of pUC18-T20 (Yu et al., 2017) in a 50 µl reaction containing 1 X Phusion HF Master Mix, 0.8 µM oligo JW151 and 0.8 µM oligo JW152, and amplicons were digested with 30 units of DpnI (New England Biolabs, Inc.) for 30 min at 37°C. The N25 promoter was amplified by PCR using 1 ng JW173 in 50 µl reactions containing 1 X Phusion HF Master Mix, 0.8 µM oligo JW179, and 0.8 µM oligo JW180, and the amplicon was isolated using a PCR purification kit (Qiagen, Inc.). Gibson assembly was performed by combining 100 ng of DpnI-digested pUC18-T20 amplicon with 25 ng of the N25 promoter amplicon in 5 µl 2 X Gibson Assembly Master Mix (New England Biolabs, Inc.) in 10 µl reactions and incubating 15 min at 50°C. The ligation mixture was diluted 2-fold with ultrapure water and was used to transform NEB 5-alpha electrocompetent cells (New England Biolabs, Inc.). Transformants were plated on LB agar plates containing 100 µg/ml carbenicillin, and recombinant plasmid DNA was isolated from individual transformants.

Plasmids pUC18-T20-N25[G_+1_G_+2_G_+3_]and pUC18-T20-N25[C_+1_C_+2_C_+3_], which were used for experiments with the N25 [G_+1_G_+2_G_+3_] promoter and the N25 [C_+1_C_+2_C_+3_] promoter, were prepared in the same manner, but using JW169 and JW170, respectively, in place of JW173

2.0 kb DNA fragments carrying single centrally located N25, N25 [G_+1_G_+2_G_+3_], or N25 [C_+1_C_+2_C_+3_] promoters were prepared by digesting plasmid pUC18-T20C2-N25-WT, plasmid pUC18-T20C2-N25 [G_+1_G_+2_G_+3_], or plasmid pUC18-T20C2-N25 [C_+1_C_+2_C_+3_] with XbaI and SbfI-HF (New England Biolabs, Inc.), followed by agarose gel electrophoresis and isolation of DNA from the gel using a gel extraction kit (Qiagen, Inc.).

Starting from the above 2.0 kb pUC18 DNA segments, constructs for single-molecule DNA nanomanipulation were prepared via ligation, at the XbaI end, to a 1.0 kb DNA fragment carrying multiple biotin residues on both strands [generated by PCR amplification of plasmid pARTaqRPOC-*lac*CONS) (Revyakin et al., 2003) with primers RPOC3140 and XbaRPOC4050 (Table S2) as described (Revyakin et al., 2003; Revyakin et al., 2004, 2005; Revyakin et al., 2006), followed by digestion with XbaI (New England Biolabs, Inc.)] and, then, at the SbfI end, to a 1.0 kb DNA fragment carrying multiple digoxigenin residues on both strands [generated by PCR amplification of plasmid pARTaqRPOC-*lac*CONS (Yu et al., 2017) with primers RPOC820 and SbfRPOC50 (Table S2) as described (Revyakin et al., 2003; Revyakin et al., 2004, 2005; Revyakin et al., 2006), followed by digestion with SbfI-HF (New England Biolabs, Inc.)]. The resulting constructs were linked to streptavidin-coated magnetic beads (1 µm diameter, MyOne Streptavidin C1; Life Technologies, Inc.), immobilized on a glass surface coated with anti-digoxigenin (prepared as in (Lionnet et al., 2012)) and calibrated as described (Revyakin et al., 2003; Revyakin et al., 2004, 2005; Revyakin et al., 2006).

#### Measurement of RNAP-dependent DNA unwinding by single-molecule DNA nanomanipulation: data collection

Experiments were performed essentially as described (Revyakin et al., 2003; Revyakin et al., 2004, 2005; Revyakin et al., 2006). We tested a 2.0 kb, mechanically extended, torsionally constrained, DNA molecule carrying N25 WT, N25 [C_+1_C_+2_C_+3_], or N25 [G_+1_G_+2_G_+3_] (superhelical density = 0.021 for experiments with positively supercoiled DNA; superhelical density = −0.021 for experiments with negatively supercoiled DNA; extension force = 0.3 pN). *E. coli* RNAP σ^70^ holoenzyme was used at 10 nM in experiments with positively supercoiled DNA and at 0.5 nM for experiments with negatively supercoiled DNA. The assay buffer contained 25 mM Na-HEPES (pH 7.9), 75 mM NaCl, 10 mM MgCl_2_, 1 mM dithiothreitol, 0.1 mg/mL BSA, and 0.1% Tween-20. NTPs (ATP, GTP, UTP, and CTP; GE Healthcare Life Sciences, Inc.) were used at 0.5 mM. Experiments were performed at 30°C. For each promoter, all data were collected on a single DNA molecule. For experiments with negatively supercoiled DNA, in which t_unwound_ > 1 h (Revyakin et al., 2003; Revyakin et al., 2004, 2005; Revyakin et al., 2006), the unwound DNA was mechanically disrupted after 1 min as described (Revyakin et al., 2003; Revyakin et al., 2004, 2005; Revyakin et al., 2006), by rotating the magnets 8 turns counterclockwise to introduce positive superhelical turns (superhelical density = 0.021), waiting for 1 min, and rotating magnets 8 turns clockwise to re-introduce negative superhelical turns (superhelical density = –0.021) and reform unwound complexes.

#### Measurement of RNAP-dependent DNA unwinding by single-molecule DNA nanomanipulation: data reduction

Raw time traces were analyzed to give DNA extension (*l*) as described (Revyakin et al., 2003; Revyakin et al., 2004, 2005; Revyakin et al., 2006). Changes in *l* resulting from DNA unwinding (Δ*l_u_*) were calculated as Δ*l_u_* = (Δ*l_obs,neg_* + Δ*l_obs,pos_*)/2, where Δ*l_obs,pos_* is the observed change in *l* in assays with positively supercoiled DNA, and Δ*l_obs,neg_* is the observed change in *l* with negatively supercoiled DNA. Extents of DNA unwinding (in base pairs) were calculated as (Δ*l_u_ /δ*) 10.4, where *δ* (55 nm) is the change in *l* per turn at the superhelical densities of this work (+/– 0.021), and 10.4 is the mean number of base pairs in each turn of B-DNA (Wang, 1979).

#### Statistical analysis

Data for single-molecule DNA nanomanipulation are means ± SEM of at least 50 technical replicates on a single, positively supercoiled DNA molecule, or at least 40 technical replicates on a single, negatively supercoiled DNA molecule.

---

**Table S1.** Data collection and refinement statistics.^a^.

| **Complex** | **RPitc,5** | **RPrtc,5 [G<sub>+1</sub>G<sub>+2</sub>G<sub>+3</sub>]** | **RPrtc,4 [C<sub>+1</sub>C<sub>+2</sub>C<sub>+3</sub>]** |
| --- | --- | --- | --- |
| **PDB code** | 7MLB | 7MLI | 7MLJ |
| **Beamline** | APS19-ID | SSRL-9-2 | SSRL-9-2 |
| **Resolution range** | 50.00 - 3.60<br>(3.66 - 3.60) | 50.00 - 3.60<br>(3.66 - 3.60) | 50.00 - 3.75<br>(3.81 - 3.75) |
| **Space group** | C 1 2 1 | C 1 2 1 | C 1 2 1 |
| **Unit cell** | 186.17 104.27<br>299.02 90 97.95 90 | 187.08 103.88 298.74<br>90 97.95 90 | 186.31 104.66 298.41<br>90 98.12 90 |
| **Unique reflections** | 66110 (3055) | 64844 (3172) | 57607 (2910) |
| **Multiplicity** | 4.0 (3.3) | 4.2 (3.9) | 4.2 (4.2) |
| **Completeness (%)** | 98.5 (92.3) | 96.84 (91.38) | 96.97 (93.60) |
| **Mean I/sigma(I)** | 10.7 | 10.8 | 14.5 |
| **Wilson B-factor** | 116.36 | 107.75 | 114.24 |
| **R-merge** | 0.116 (0.845) | 0.151 (1.36) | 0.103 (1.031) |
| **CC1/2** | (0.624) | (0.912) | (0.837) |
| **Reflections used in refinement** | 65151 (6097) | 64469 (5989) | 57263 (5466) |
| **R-work/R-free** | 0.2330/0.2697 | 0.2432/0.2918 | 0.2654/0.2833 |
| **Number of non-H atoms** | 28400 | 28502 | 28489 |
| **RMS(bonds)** | 0.002 | 0.002 | 0.002 |
| **RMS(angles)** | 0.53 | 0.54 | 0.52 |
| **Ramachandran favored (%)** | 98.44 | 98.30 | 97.84 |
| **Ramachandran outliers (%)** | 0.09 | 0.14 | 0.12 |
| **Rotamer outliers (%)** | 6.37 | 6.37 | 5.63 |
| **Clashscore** | 6.16 | 6.41 | 6.22 |

<sup>a</sup> Statistics for the highest-resolution shell are shown in parentheses.

**Table S2.**
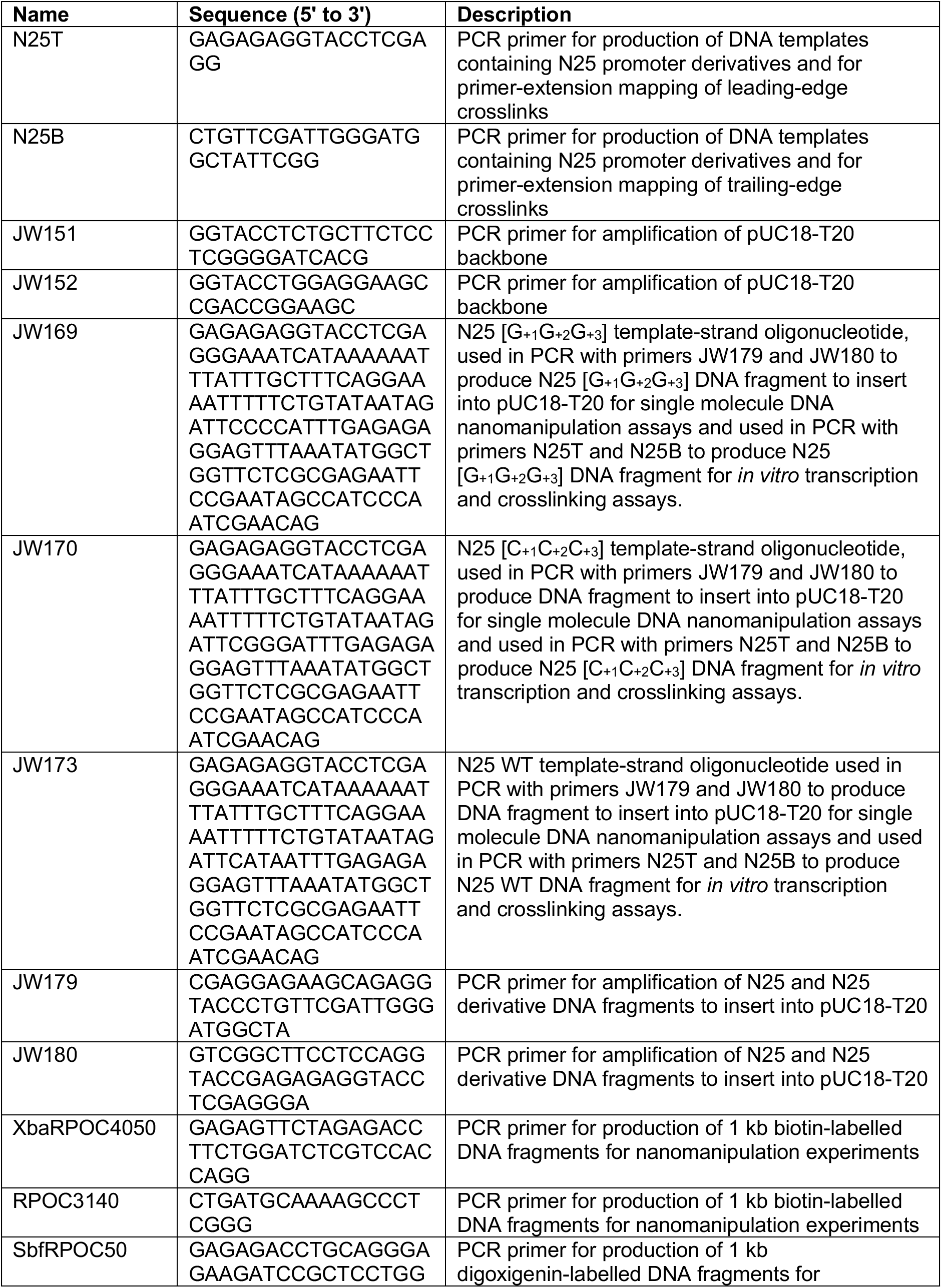
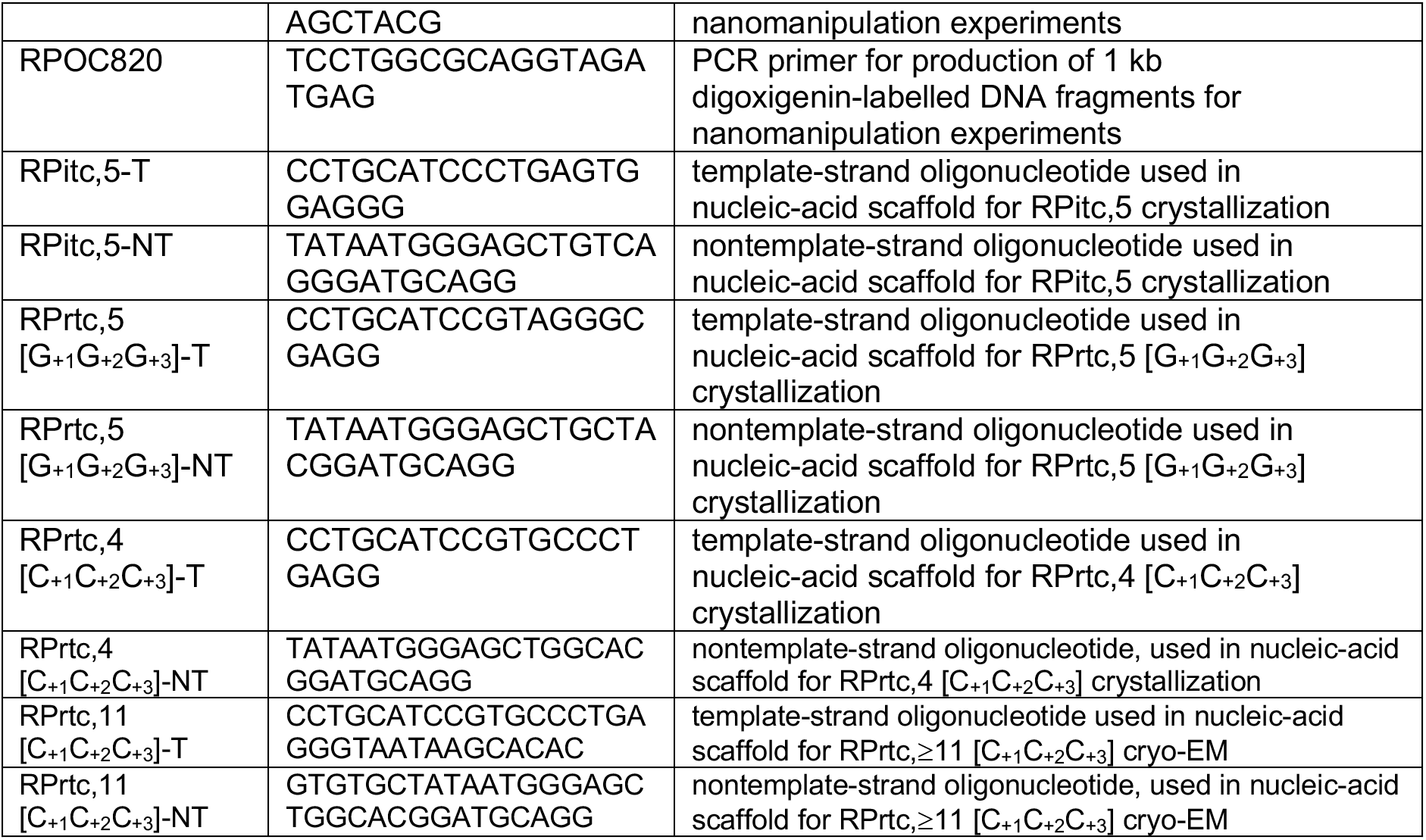
Oligonucleotides.

## Supplemental Figure Legends

**Figure S1.**
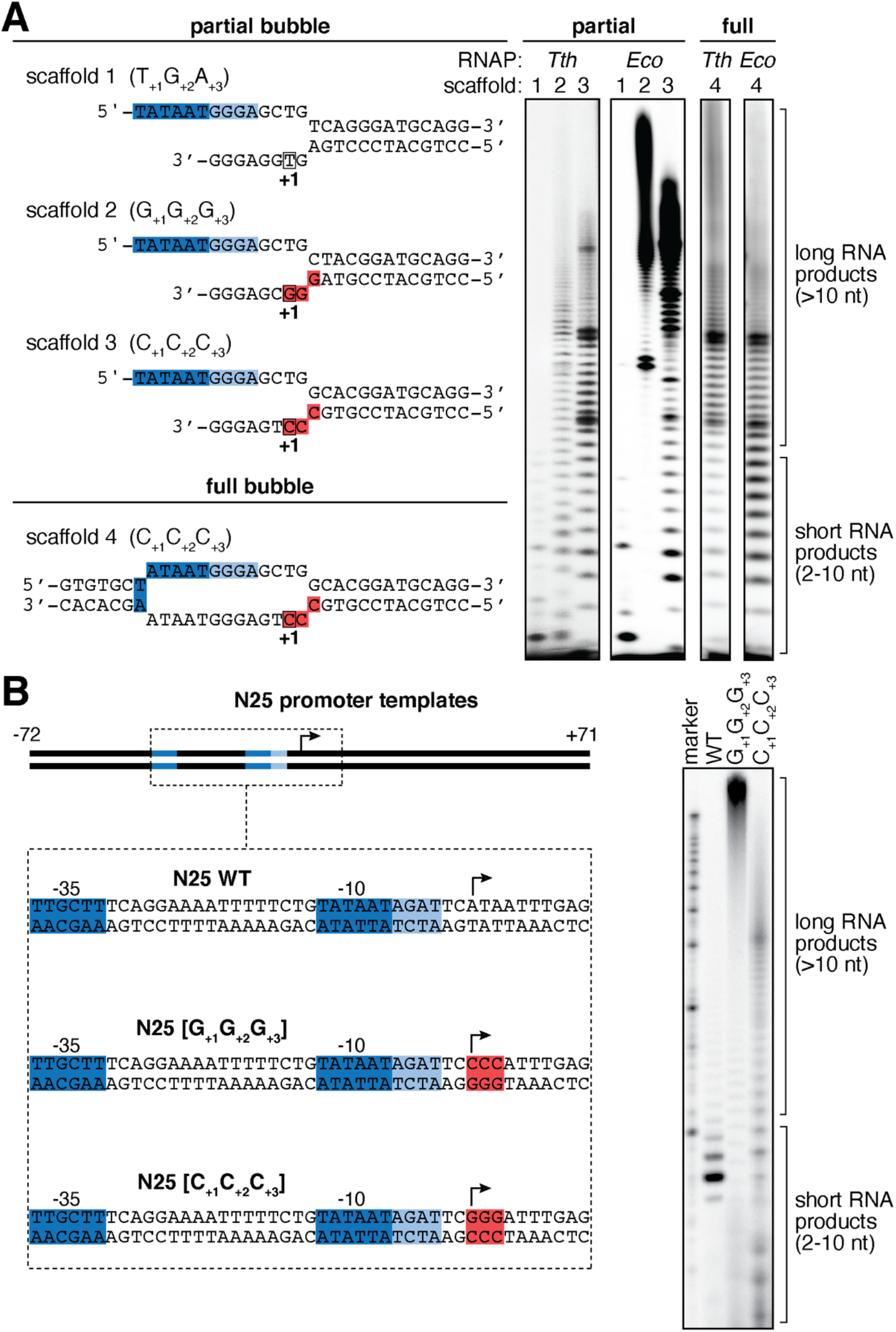
Standard transcription initiation and reiterative transcription initiation. **(A)** *In vitro* transcription experiments with nucleic-acid scaffolds used in crystal structure determination of Figures 1-2 (“partial bubble”) and cryo-EM structure determination of Figures 3-4 (“full bubble”). *Left*, nucleic-acid scaffolds (promoter −35 and −10 elements, blue; promoter discriminator elements, light blue; template-strand homopolymeric sequences, red; non-complementary sequences, raised and lowered nucleotides; transcription start sites, +1). *Right*, *in vitro* transcription products with *T. thermophilus* RNAP σ^A^ holoenzyme (*Tth*) and *E. coli* RNAP σ^70^ holoenzyme (*Eco*) for partial-bubble scaffold 1 (template-strand non-homopolymeric transcription-start-site sequence T_+1_G_+1_A_+1_; transcription with ATP and UTP; standard transcription initiation), partial-bubble scaffold 2 (template-strand homopolymeric sequence G_+1_G_+2_G_+3_; transcription with CTP; reiterative transcription initiation), partial-bubble scaffold 3 (template-strand homopolymeric sequence C_+1_C_+2_C_+3_; transcription with GTP; reiterative transcription initiation), and full-bubble scaffold 4 (template-strand homopolymeric sequence C_+1_C_+2_C_+3_; transcription with GTP; reiterative transcription initiation). **(B)** *In vitro* transcription experiments with bacteriophage T5 N25 promoter derivatives used in photocrosslinking and single-molecule DNA-nanomanipulation experiments of Figures 5-6. *Left*, promoter sequences (colors as in panel A). *Right*, *in vitro* transcription products with and *E. coli* RNAP σ^70^ holoenzyme (*Eco*) for bacteriophage N25 WT (transcription with ATP and UTP; standard transcription initiation), N25 [G_+1_G_+2_G_+3_] (template-strand homopolymeric sequence G_+1_G_+2_G_+3_; transcription with CTP; reiterative transcription initiation), and N25 [C_+1_C_+2_C_+3_] (template-strand homopolymeric sequence C_+1_C_+2_C_+3_; transcription with GTP; reiterative transcription initiation). RNA products 2-10 nt in length are products of standard transcription initiation or reiterative transcription initiation; RNA products >10 nt in length are products of reiterative transcription initiation.

**Figure S2.**
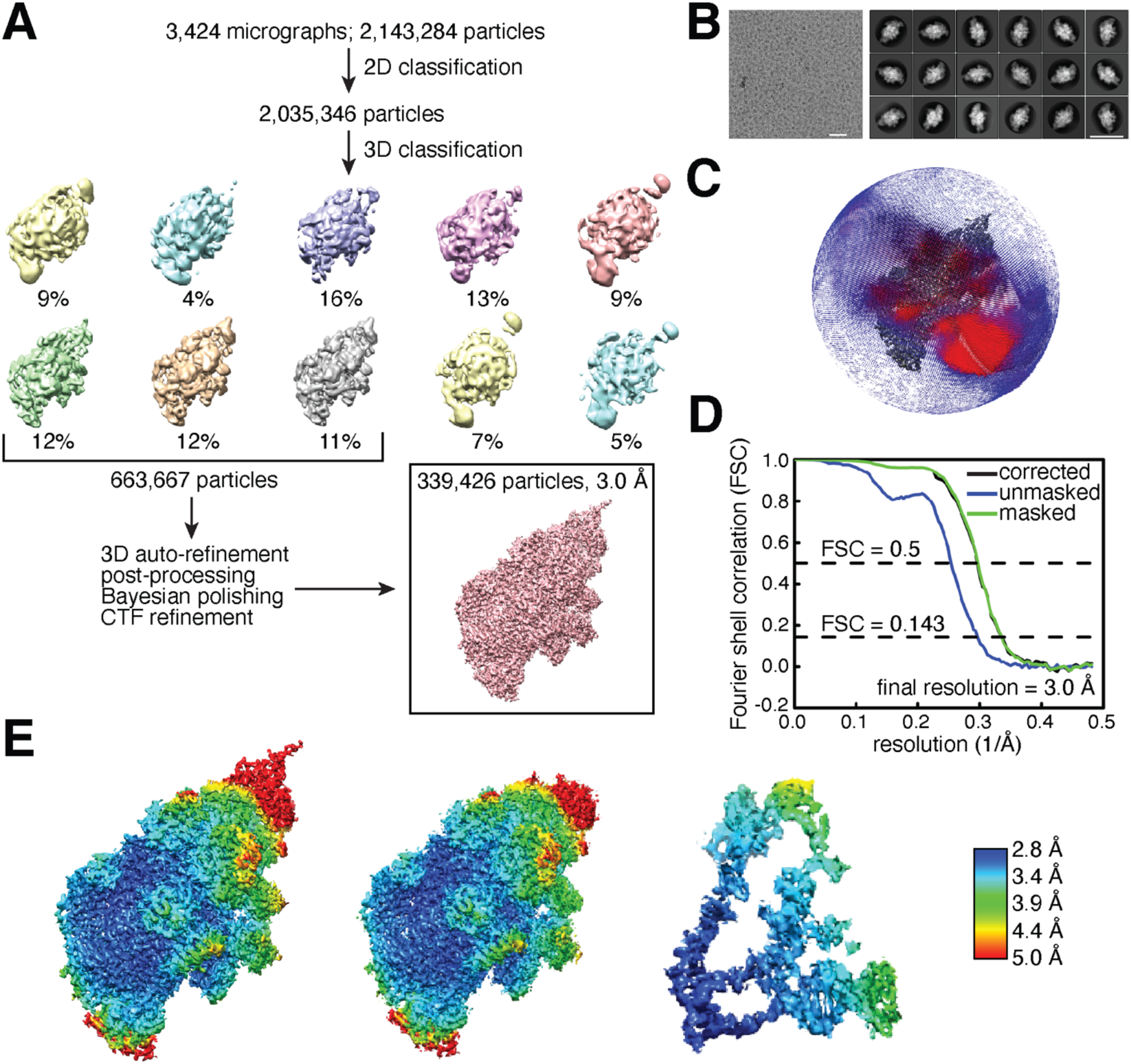
Cryo-EM structure determination: RPrtc,≥11 [C_+1_C_+2_C_+3_]. **(A)** Data processing scheme. **(B)** Representative electron micrograph (left; 50 nm scale bar) and representative class averages (right, 20 nm scale bar). **(C)** Orientational distribution. **(D)** Fourier-shell-correlation (FSC) plot. **(E)** EM density maps colored by local resolution. Left, overall structure. Center, overall structure, omitting β’ non-conserved region. Right, DNA and RNA in structure. View orientations as in Figures 3A-B, D.

**Figure S3.**
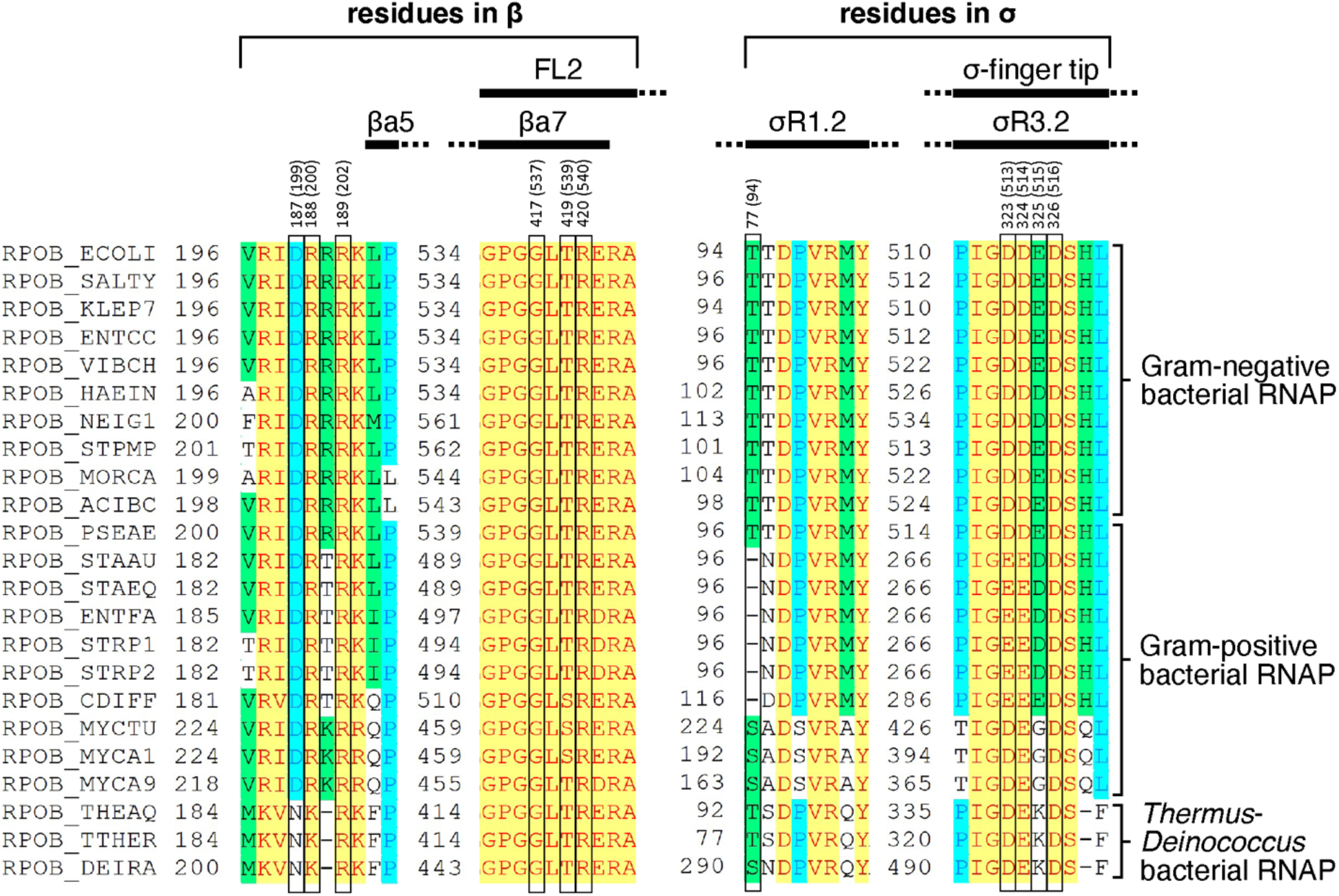
RNAP and σ residues that make protein-RNA interactions in alternative RNA pathway and alternative RNA exit: sequence-conservation patterns of. Locations of residues that make protein-RNA interactions with rN6-rN11 in the structure of RPrtc,≥11 [C_+1_C_+2_C_+3_] in the sequences of RNAP β subunit (left) and σ (right). Sequence alignments for β and σ^70^/σ^A^ for Gram-negative, Gram-positive, and *Thermus*-*Deinococcus*-clade bacterial species showing locations of RNAP residues that contact protein-RNA interactions with rN6-rN11 (black rectangles; numbered as in *T. thermophilus* and, in parentheses, as in *E. coli*; identities from Figure 4C), and locations of RNAP and σ structural elements and conserved regions (black bars; Feklistov et al., 2014; Lane and Darst, 2010). Species are as follows: *E. coli* (ECOLI), *Salmonella typhimurium* (SALTY), *Klebsiella pneumoniae* (KLEP7), *Enterobacter cloacae* (ENTCC), *Vibrio cholerae* (VIBCH), *Haemophilus influenzae* (HAEIN), *Neisseria gonorrhoeae* (NEIG1), *Stenotrophomonas maltophilia* (STPMP), *Moraxella catarrhalis* (MORCA), *Acinetobacter baumannii* (ACIBC), *Pseudomonas aeruginosa* (PSEAE), *Staphylococcus aureus* (STAAU), *Staphylococcus epidermidis* (STAEQ), *Enterococcus faecalis* (ENTFA), *Streptococcus pyogenes* (STRP1), *Streptococcus pneumoniae* (STRP2), *Clostridium difficile* (CDIFF), *Mycobacterium tuberculosis* (MYCTU), *Mycobacterium avium* (MYCA1), *Mycobacterium abscessus* (MYCA9), *Thermus aquaticus* (THEAQ), and *Thermus thermophilus* (TTHER), and *Deinococcus radiodurans* (DEIRA).

**Figure S4.**
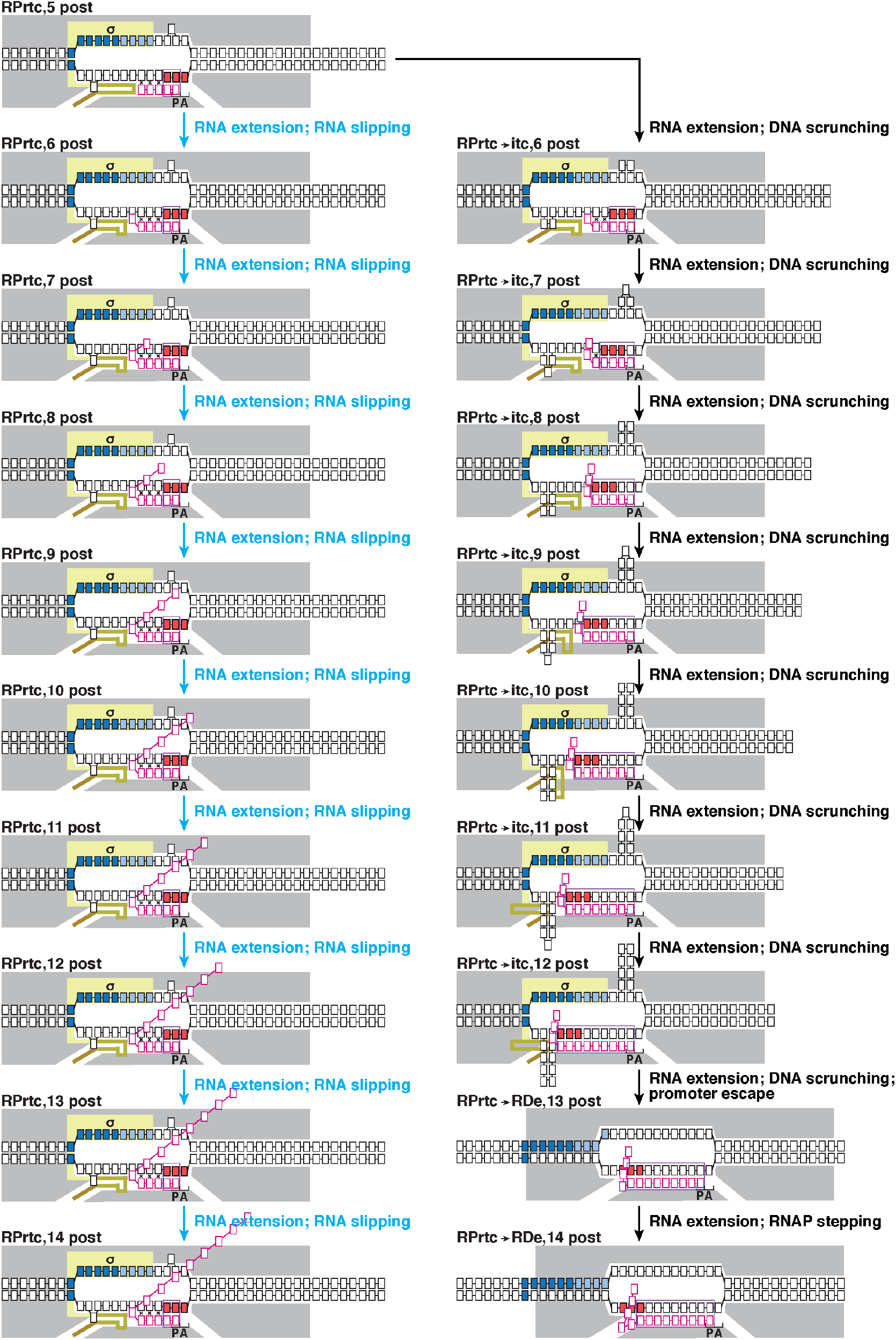
Hypothesized mechanisms for synthesis of long RNA products ( >10 nt) through reiterative transcription initiation. The part of the σ finger that occupies the RNAP hybrid binding site is colored dark yellow; the part of the σ region-3/region-4 linker that occupies the RNAP RNA-exit channel is colored brown. Non-templated, reiterative-transcription-dependent RNA nucleotides are colored magenta; templated, standard-transcription-dependent RNA nucleotides are colored orange. Other symbols and colors are as in Figures 5 and 7. **(Left)** Hypothesized mechanism for synthesis of long RNA products comprising only non-templated, reiterative-transcription-dependent nucleotides (“non-productive” reiterative transcription initiation by RPrtc, without displacement of the σ finger, displacement of the σ region-3/region-4 linker, and promoter escape). **(Right)** Hypothesized mechanism for synthesis of long RNA products comprising non-templated, reiterative-transcription-dependent nucleotides at their 5’ ends followed by templated, standard-transcription-dependent nucleotides (“productive” reiterative transcription initiation, entailing switching from reiterative transcription initiation by RPitc to standard transcription initiation by RPrtc**→**itc, resulting in displacement of the σ finger, displacement of the σ region-3/region-4 linker, promoter escape, optional σ release, and transcription elongation using an RNAP-stepping mechanism).

